# Time dependent changes in feeding behavior and energy balance associated with weight gain in mice fed obesogenic diets

**DOI:** 10.1101/2024.01.10.575043

**Authors:** Payam A. Fathi, Michelle B. Bales, Julio E. Ayala

## Abstract

Obesity is characterized by dysregulated homeostatic mechanisms resulting in positive energy balance, yet when this dysregulation occurs is unknown. We assessed the time course of alterations to behaviors promoting weight gain in male and female mice switched to obesogenic 60% or 45% high fat diet (HFD). Switching mice to obesogenic diets promotes transient bouts of hyperphagia during the first 2 weeks followed by persistent caloric hyperphagia. Energy expenditure increases but not sufficiently to offset increased caloric intake, resulting in a sustained net positive energy balance. Hyperphagia is associated with consumption of calorically larger meals (impaired satiation) more frequently (impaired satiety) particularly during the light-cycle. Running wheel exercise delays weight gain in 60% HFD-fed male mice by enhancing satiation and increasing energy expenditure. However, exercise effects on satiation are no longer apparent after 2 weeks, coinciding with weight gain. Thus, exposure to obesogenic diets engages homeostatic regulatory mechanisms for ∼2 weeks that ultimately fail, and consequent weight gain is characterized by impaired satiation and satiety. Insights into the etiology of obesity can be obtained by investigating changes to satiation and satiety mechanisms during the initial ∼2 weeks of HFD exposure.

**What is already known about this subject?:**

- Obesity is associated with dysregulated homeostatic mechanisms.
- Increased caloric consumption contributes to obesity.
- Obese rodents tend to eat larger, more frequent meals.

**What are the new findings in your manuscript?:**

- Exposure to obesogenic diets promotes transient attempts to maintain weight homeostasis.
- After ∼2 weeks, caloric hyperphagia exceeds increased energy expenditure, promoting weight gain.
- This is associated with consumption of larger, more frequent meals.

**How might your results change the direction of research or the focus of clinical practice?:** Our findings suggest that molecular studies focusing on mechanisms that regulate meal size and frequency, particularly those engaged during the first ∼2 weeks of obesogenic diet feeding that eventually fail, can provide unique insight into the etiology of obesity.

## Introduction

Obesity results from caloric intake exceeding energy expenditure (EE). A general model for body weight regulation posits that weight gain triggers feedback mechanisms primarily in the central nervous system to subsequently decrease caloric intake, increase EE, and consequently reduce body weight (1). Obesity has been characterized as being associated with defects in these feedback mechanisms. However, evidence that these mechanisms are dysregulated has primarily been obtained comparing lean and obese subjects (mice, rats, humans). This is potentially problematic because homeostatic mechanisms may be preserved in obesity to defend elevated body weight (2, 3). Therefore, characterizing changes in parameters that regulate body weight prior to obesity onset can provide insights into potential mechanisms associated with excess weight gain.

Studies in rats show exposure to a high-fat diet (HFD) promotes a large, yet transient increase in caloric intake followed by sustained, slightly elevated daily caloric intake characterized by increased meal size and meal rate by calories but decreased meal number (4–7). These phenotypes are relevant since rats prone to greater weight gain consume more calories on a per meal basis without altering meal numbers (4, 7). Mice show a similar pattern characterized by a transient increase in caloric intake when switched to a HFD followed by a progressive decrease in calories consumed (8–11). A limitation of these studies is that HFD exposure was brief (3-7 days), and the primary endpoint for consumption was gross food intake. More nuanced factors that could influence weight gain, such as meal patterns, were not assessed. The importance of this limitation is highlighted by the fact that genetic mouse models of obesity display increased meal size without increased daily caloric intake (12–15).

Sex and timing of exposure to obesogenic diets also affect weight gain. Male rodents increase caloric intake and body weight more than female rodents (16, 17). Mice fed obesogenic diets only during the light phase gain more weight than mice fed the same diet only during the dark phase (18). Access to HFD rapidly disrupts circadian food intake patterns, and mice no longer consume the majority of meals during the dark cycle (19).

Taken together, factors such as meal patterns, sex, and timing of food intake influence weight gain associated with obesogenic diets. When these factors are altered during the progression towards obesity is unknown. The present studies investigated the time scale and magnitude of changes to these components during the progression towards obesity. Our findings demonstrate that upon exposure to HFD, behaviors and metabolic phenotypes engaged to prevent weight gain rapidly become dysregulated. Furthermore, the magnitude of these changes is influenced by the type of obesogenic diet, sex, and light vs. dark cycles. This provides a blueprint for investigating molecular mechanisms whose dysregulation contributes to developing obesity.

## Methods

### Animals

Male and female C57Bl6/J mice were studied at 10-12 weeks of age. In-house breeders were fed “breeder” diet (5LJ5 – PicoLab Mouse Diet, LabDiet) and offspring were placed on chow (5L0D – PicoLab Rodent Diet) at weaning (21 days). Experiments were conducted within an 18-month period in offspring produced by multiple simultaneous breeders from the same founders to minimize genetic drift. Mice were maintained on a 12-12 light-dark cycle 0600–1800) at 23°C. Experimental procedures were approved by the Institutional Animal Care and Use Committee at Vanderbilt University.

### Diet Switch and Running Wheel Experiments

Mice were single housed in Promethion Metabolic chambers (Sable Systems International) with ad-lib access to chow (2.86 kCal/gram; PicoLab Rodent Diet 5L0D) for 5-7 days. Mice were then switched to 60% HFD (5.24 kCal/gram; Research Diets Inc, D12492) for 4 weeks or 45% HFD (4.73 kCal/gram; Research Diets Inc. D12451) for 3 weeks. A separate cohort was switched to 60% HFD or remained on chow for 3 weeks. For wheel running experiments, single-housed male mice had access to chow and locked running wheels for 5 days. Mice were then switched to 60% HFD and were randomly selected to remain with running wheels locked or have running wheels unlocked for 4 weeks. External interventions (e.g., adding food/water, changing cages) are indicated in each figure.

### Metabolic Measurements

Food intake, water intake, locomotor activity, respiratory exchange ratio, and EE were continuously measured across five-minute intervals. Energy balance was determined by subtracting total EE from energy intake across 12h light-dark cycles. Total EE for each light-dark cycle was calculated as the average EE over the 12h time interval multiplied by 12. Energy intake was calculated by multiplying the mass of food consumed over the corresponding 12h-intervals with the diet caloric density. A technical failure eliminated locomotor activity data after week 1 in 60% HFD-fed females. Body weights were measured weekly and body composition before and after the HFD period was obtained by NMR (Bruker Optics).

### Meal Parameters

A meal was defined as food removal spanning at least 30 seconds, a minimum of 10 milligrams to account for inherent sensor limitations, a maximum of 500 milligrams to account for spillage, and a threshold of 5 minutes for inter-meal intervals based on prior rodent literature (20–23). 60% HFD was packed in the hopper to reduce spillage. Cages were inspected daily for excessive spillage (food on cage bottom) and noted. If total food intake and meal number for a given day was greater than 1.5x standard deviation of the mean for its respective feeding period, then this was considered excessive spillage, and intake and meal pattern data for that day were excluded. Excessive spillage was only observed with the 45% HFD experiments.

### Meal Type Analysis

Quartile distributions for meal types were defined by assembling meal sizes from all chow-fed male mice to determine the following meal types: Type I: 10 – <60 mg; Type II: 60 – <160 mg; Type III: 160 – <280 mg; and Type IV: >280 mg. Meal types were compared by determining the weekly proportion of a given meal type within each animal across each week. Parts of whole analysis used compiled averages of meal types across sex and diet groups. Histograms were generated using bin sizes of 20 mg or .2 kCal and curve-fit using non-linear regression.

### Statistical Analysis

Raw data were processed using the Sable Systems Macro Interpreter (v23.6). Graphs were generated and data statistically analyzed using repeated measures one-way ANOVA and Tukey’s post-hoc or two-way ANOVA with the Sidak’s post-hoc as necessary using GraphPad Prism (v10). Significance was set at p<0.05. Figures were assembled using Adobe Illustrator (v28.0).

## Results

### Metabolic phenotypes in mice switched to 60% HFD

When switching male mice to 60% HFD, food intake by mass is higher at week 4 of 60% HFD (**Figure 1A**), with a tendency for higher light cycle mass consumption throughout the HFD period (**Supplemental Figure 1A-B**). Food intake by mass is unaffected in female mice switched to 60% HFD (**Figure 1D**) during both cycles (**Supplemental Figure 1C-D**). Accounting for caloric density, both sexes display two cycles of rapid increases and decreases in calories consumed during the first 2 weeks of 60% HFD exposure followed by a sustained elevation in daily caloric intake (**Figure 1B, 1E**). This occurs during both light and dark cycles (**Supplemental Figure 1E-H**). This pattern does not correspond with interventions such as restocking of food or cage changes.

**Figure 1.**
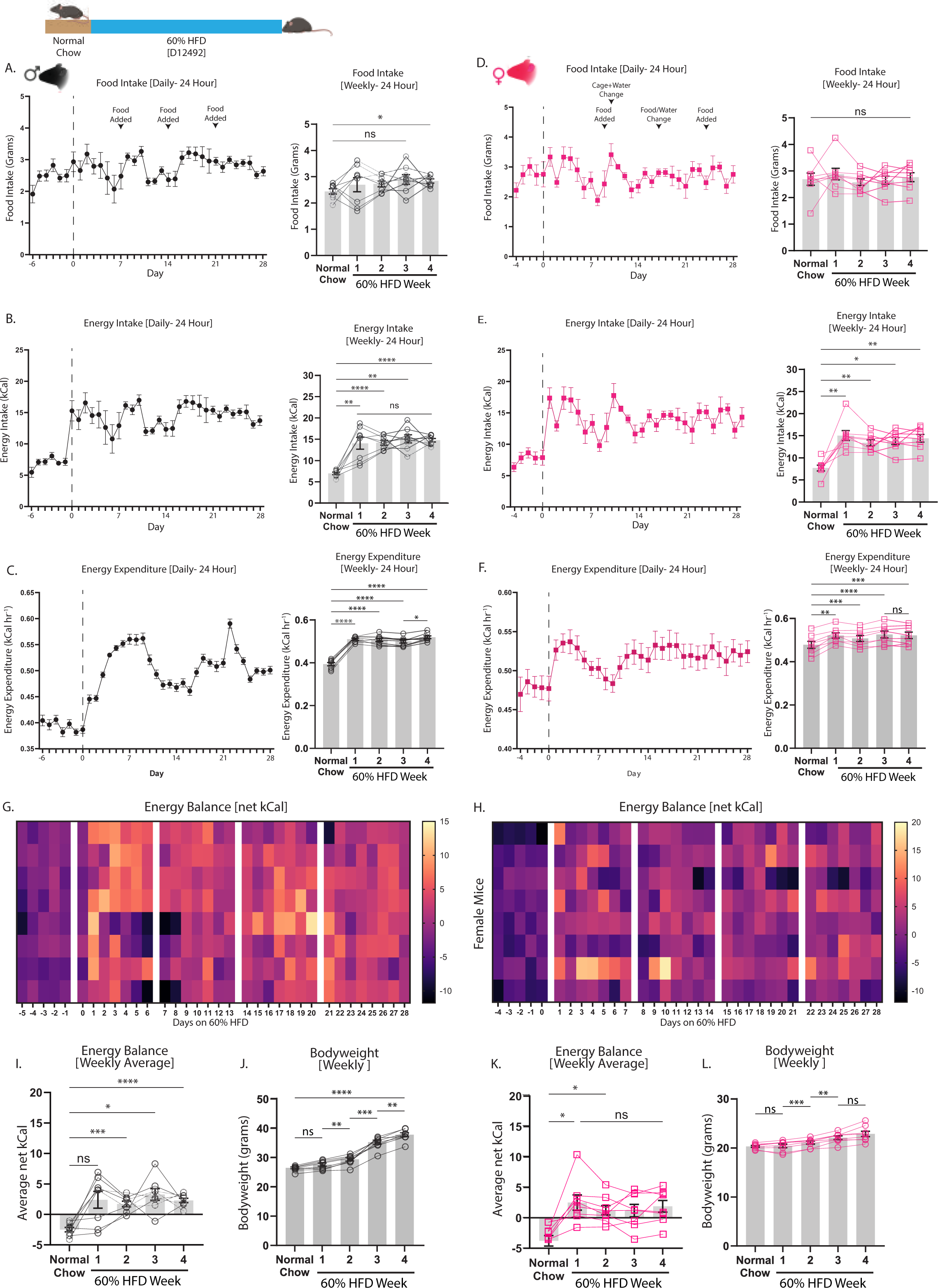
Food intake and energy balance in mice switched to 60% HFD. Mice were placed in metabolic chambers and were fed with chow for up to 6 days. Mice were then switched to 60% HFD at day 0 (dashed line) and remained on 60% HFD for 28 days. (A-C, G, I, J) represent data from male mice (black) and (D-F, H, K, L) represent data from female mice (pink). (A, D) Daily and average weekly food intake by mass (grams). (B, E) Daily and average weekly caloric intake. (C, F) Daily and average weekly energy expenditure. (G, H) Heat map showing shifts in net daily energy balance. (I, K) Average weekly energy balance. (J, L) Average weekly body weight. Arrows indicate external interventions such as adding food/water or cage changes. Data are shown as mean ± SEM for N=8 per sex and were statistically analyzed using a repeated measures one-way ANOVA with Tukey multiple comparison tests. P-value definitions: not-significant (ns) >0.05, * <0.05, **<0.01, ***<0.001, **** <0.00001.

Switching to 60% HFD increases EE in males and females (**Figure 1C, 1F**) during both light and dark cycles (**Supplemental Figure 1I-L**). Accounting for changing body weight, analysis of covariance (ANCOVA) shows EE remains higher during the HFD period in males and trends higher in females (**Supplemental Figure 2A**). Daily EE follows a similar pattern as caloric intake, characterized by transient increases upon switching to the HFD followed by a sustained increase after 2 weeks of the diet switch (**Figure 1C, 1F**). Male and female mice display sustained positive energy balance and increasing body weight starting on week 2 of HFD exposure (**Figure 1G-L**). Lean and fat mass increase in 60% HFD-fed male mice and tend to increase in female mice (**Supplemental Figure 2B**).

Respiratory exchange ratio (RER) decreases upon exposure to 60% HFD in both sexes (**Supplemental Figure 2C-D**), most robustly during the dark cycle (**Supplemental Figure 1M-P**). Switching to 60% HFD rapidly decreases water intake and locomotor activity in both sexes (**Supplemental Figure 2E-H**) primarily due to changes during the dark cycle (**Supplemental Figure 1Q-W**).

### Meal patterns in mice switched to 60% HFD

Meal numbers rapidly increase in both sexes and are sustained throughout the 60% HFD period (**Figure 2A-B**) in both light and dark cycles (**Supplemental Figure 3A-D**). Inter-meal interval and meal duration decreases in both sexes (**Figure 2C-F**), with robust effects on inter-meal interval during the light cycle (**Supplemental Figure 3E-H**) and meal duration during both cycles (**Supplemental Figure 3I-L**). Meal size by mass decreases in male and female mice (**Figure 2G-H**) primarily during the dark cycle (**Supplemental Figure 3M-P**). Accounting for caloric density, meal size significantly increases in male mice after the second week of 60% HFD exposure but only trends higher in female mice (**Figure 2I-J**). Male mice primarily increase meal caloric content during the light cycle (**Supplemental Figure 3Q-R**), and female mice do so during the dark cycle (**Supplemental Figure 3S-T**). There are transient increases and attempts to restore meal size during the first two weeks of 60% HFD exposure. Rates of meal consumption by mass or caloric content increase upon switching to 60% HFD in both males and females (**Figure 2K-N**) primarily during the light cycle (**Supplemental Figure 3U-AB**).

**Figure 2.**
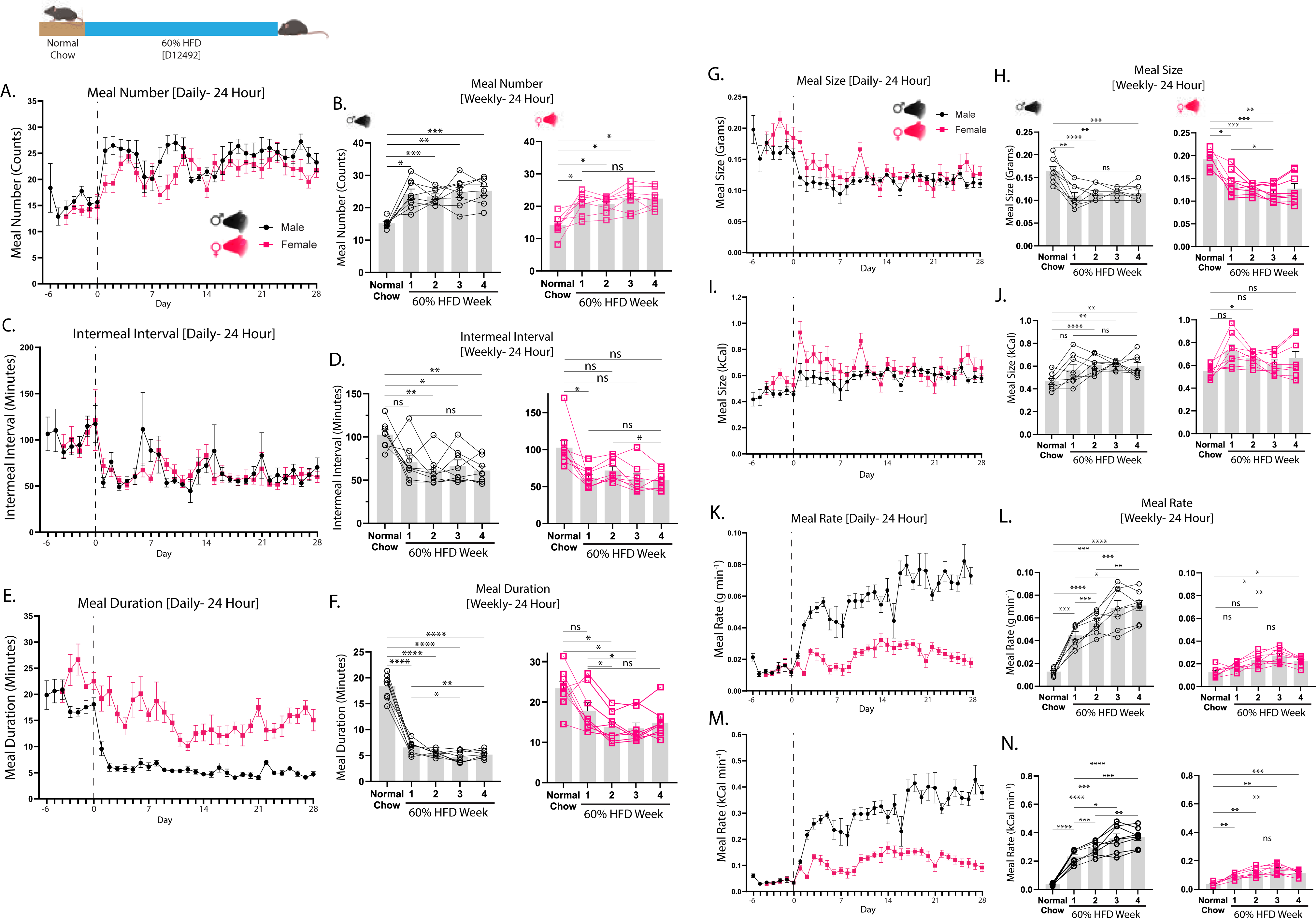
Feeding behaviors in mice switched to 60% HFD. Feeding behavior analysis in male mice (black) and female mice (pink) switched from chow to 60% HFD in metabolic chambers at day 0 (dashed line). (A) Daily and (B) average weekly meal numbers. (C) Daily and (D) average weekly intermeal intervals. (E) Daily and (F) average weekly meal duration. (G) Daily and (H) average weekly meal size by mass. (I) Daily and (J) average weekly meal size by calorie. (K) Daily and (L) average weekly meal rate in grams consumed per minute. (M) Daily and (N) average weekly meal rate in calories consumed per minute. Data are shown as mean ± SEM for N=8 per sex and were statistically analyzed using a repeated measures one-way ANOVA with Tukey multiple comparison tests. P-value definitions: not-significant (ns) >0.05, * <0.05, **<0.01, ***<0.001, **** <0.00001.

Changes to metabolic and meal parameters in male mice switched to 60% HFD are not observed in mice remaining on chow for 3 weeks (**Supplemental Figure 4**). Furthermore, male mice gain more weight even though they increase EE more robustly than female mice (**Supplemental Figure 5C-D**). Male mice display a greater increase in meal rates than females (**Supplemental Figure 5I-J**).

### Metabolic phenotypes in mice switched to 45% HFD

When switched to 45% HFD, food intake by mass transiently increases in both sexes (**Figure 3A, 3D**) during light and dark cycles (**Supplemental Figure 6A-D**). Caloric intake increases during weeks 1 and 3 (**Figure 3B, 3E**), and there is a tendency for this to occur during both light and dark cycles (**Supplemental Figure 6E-H**). Both sexes rapidly increase EE (**Figure 3C, 3F**) during the light and dark cycles (**Supplemental Figure 6I-L**). ANCOVA accounting for body weight changes shows EE significantly increases in male mice and trends higher in female mice (**Supplemental Figure 7A**). Energy balance becomes more positive and body weights gradually increase in both sexes after switching to 45% HFD (**Figure 3G-L**). Lean and fat mass increase in 45% HFD-fed male mice but only lean mass increases in female mice (**Supplemental Figure 7B**).

**Figure 3.**
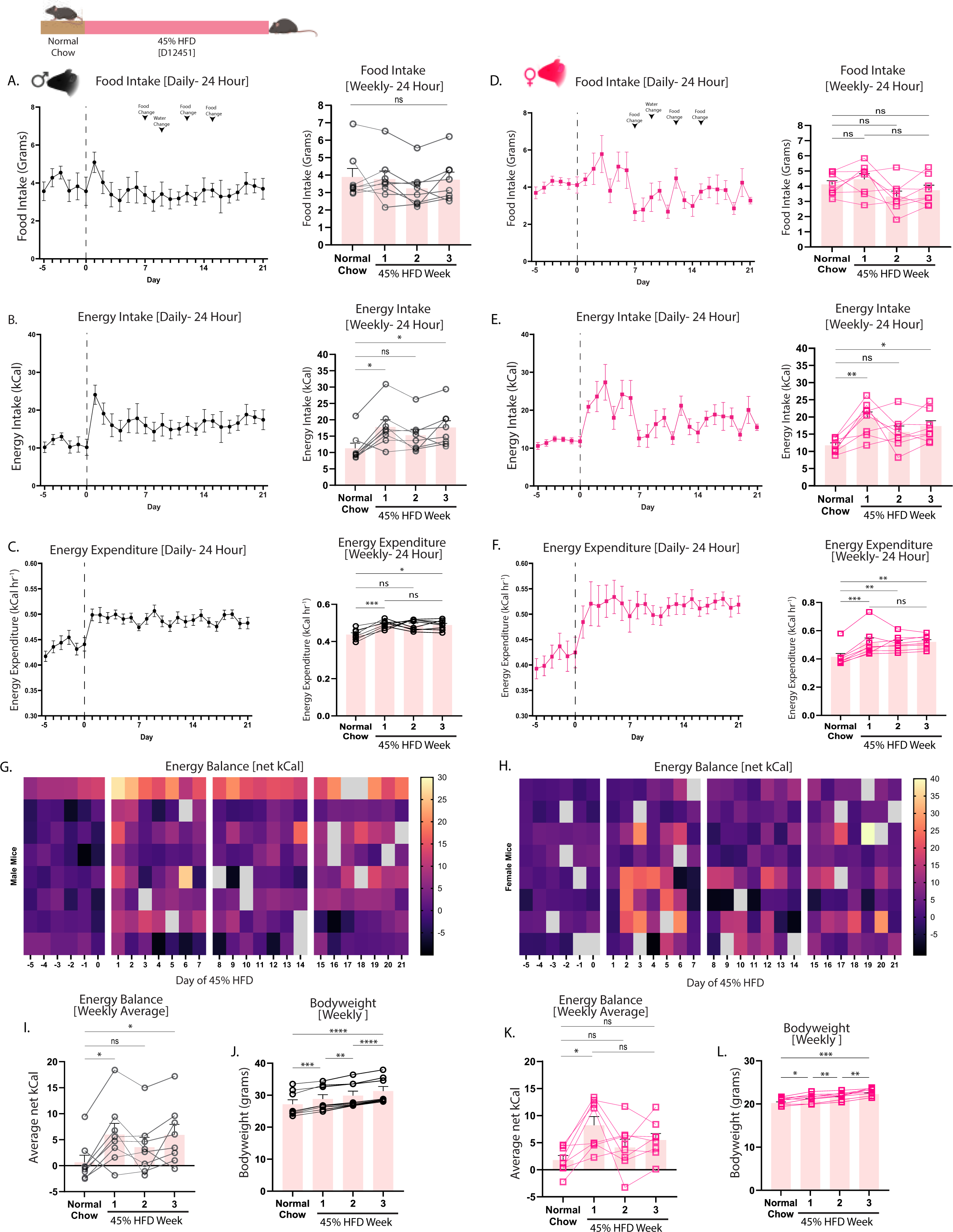
Food intake and energy balance in male and female mice switched to 45% HFD. Mice were placed in metabolic chambers and were fed with chow for 5 days. Mice were then switched to 45% HFD at day 0 (dashed line) and remained on 45% HFD for 21 days. (A-C, G, I, J) represent data from male mice (black) and (D-F, H, K, L) represent data from female mice (pink). (A, D) Daily and average weekly food intake by mass (grams). (B, E) Daily and average weekly caloric intake. (C, F) Daily and average weekly energy expenditure. (G, H) Heat map showing shifts in net daily energy balance. Blank squares indicate days in which data were excluded due to excessive spillage. (I, K) Average weekly energy balance. (J, L) Average weekly body weight. Arrows indicate external interventions such as adding food/water or cage changes. Data are shown as mean ± SEM for N=8 per sex and were statistically analyzed using a repeated measures one-way ANOVA with Tukey multiple comparison tests. P-value definitions: not-significant (ns) >0.05, * <0.05, **<0.01, ***<0.001, **** <0.00001.

Switching to 45% HFD decreases RER and water intake in both males and females, and there is only a significant activity decrease in males (**Supplemental Figure 7C-M**). These decreases are observed almost exclusively during the dark cycle (**Supplemental Figure 6M-X**).

### Meal patterns in mice switched to 45% HFD

Mice switched to 45% HFD display no significant change in meal number or inter-meal interval in either sex (**Figure 4A-D**) during both light and dark cycles (**Supplemental Figure 8A-D**). Meal duration significantly decreases in both sexes (**Figure 4E-F**) during light and dark cycles (**Supplemental Figure 8I-L**). Meal size by mass is unaffected in male mice and tends to decrease in female mice (**Figure 4G-H**) primarily due to effects during the light cycle (**Supplemental Figure 8M-P**). Accounting for caloric density, caloric intake per meal increases in male mice, primarily during the dark cycle, but remains unaltered in female mice (**Figure 4I-J**, **Supplemental Figure 8Q-T**). Meal rates by mass and caloric content increase in both sexes and during both light and dark cycles (**Figure 4M-N, Supplemental Figure 8U-AB**). Mice switched to 45% HFD display significant food spillage, so intake and meal patterns likely overestimate actual intake and contribute to increased data variability.

**Figure 4.**
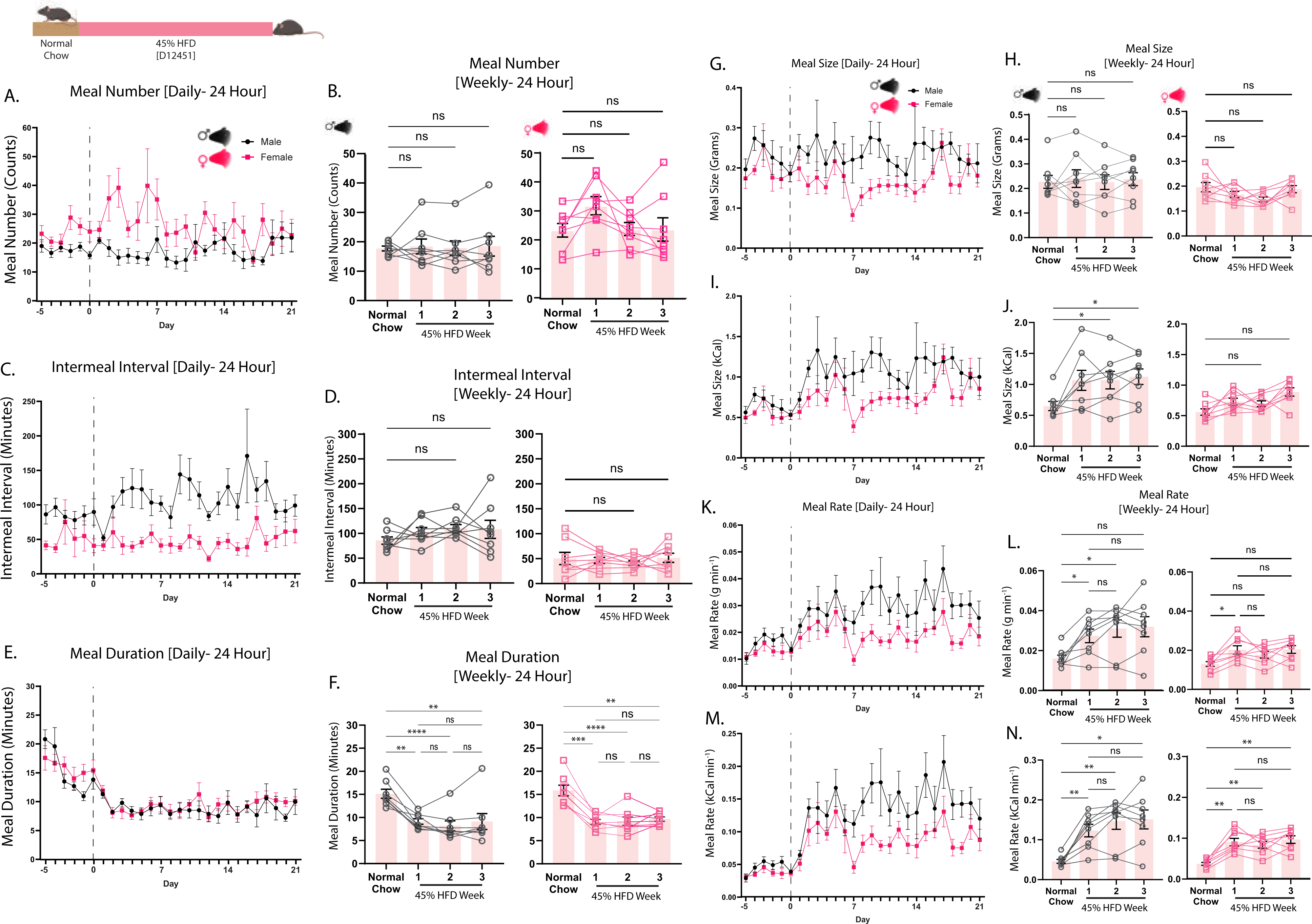
Feeding behaviors in mice switched to 45% HFD. Feeding behavior analysis in male mice (black) and female mice (pink) switched from chow to 45% HFD in metabolic chambers at day 0 (dashed line). (A) Daily and (B) average weekly meal numbers. (C) Daily and (D) average weekly intermeal intervals. (E) Daily and (F) average weekly meal duration. (G) Daily and (H) average weekly meal size by mass. (I) Daily and (J) average weekly meal size by calorie. (K) Daily and (L) average weekly meal rate in grams consumed per minute. (M) Daily and (N) average weekly meal rate in calories consumed per minute. Data are shown as mean ± SEM for N=8 per sex and were statistically analyzed using a repeated measures one-way ANOVA with Tukey multiple comparison tests. P-value definitions: not-significant (ns) >0.05, * <0.05, **<0.01, ***<0.001, **** <0.00001.

Females gain less weight than males when switched to 45% HFD, and this is associated with a greater increase in EE in females (**Supplemental Figure 9C-D**). Males display greater increases in meal rates, significantly so at 2 weeks of 45% HFD exposure (**Supplemental Figure 9I-J**).

### Meal-typing in mice switched to HFD

Meals of very low mass and caloric content comprise the largest set of meals consumed during the chow period (**Figure 5A-D**). When switched to 60% HFD, male mice decrease consumption of meals of very low and larger masses (**Figure 5A**), resulting in a rightward shift towards a larger proportion of meals with higher caloric content (**Figure 5B**). Female mice switched to 60% HFD decrease the proportion of large meals and increase the proportion of small meals consumed (**Figure 5C**), but the proportion of meals with higher caloric content increases (**Figure 5D**). When switched to 45% HFD, there is a leftward shift in meal mass histograms in both male and female mice with a corresponding flattening of histogram spread in meal size by calories (**Figure 5E-H**).

**Figure 5.**
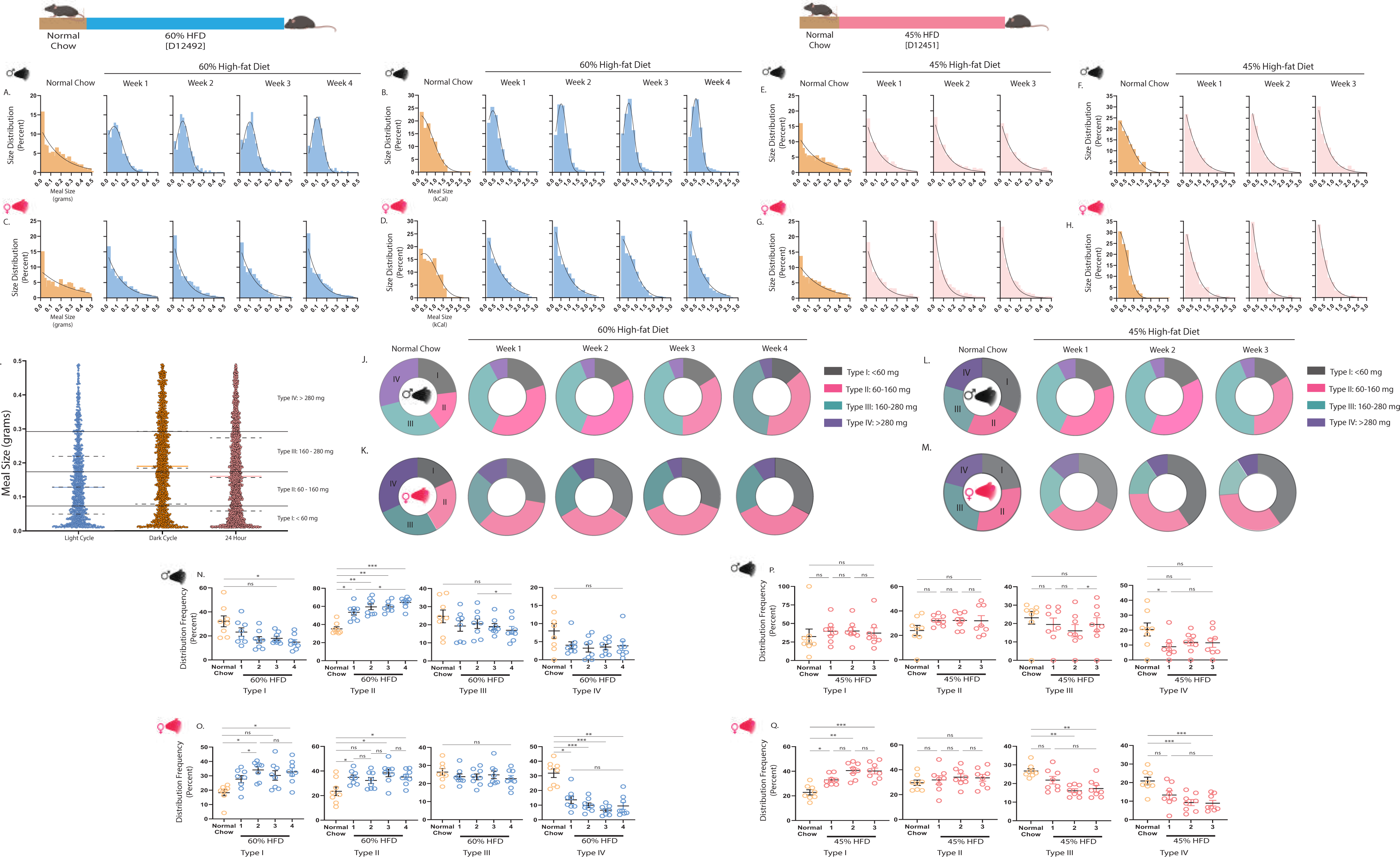
Meal patterns of male and female mice fed obesogenic diets. Meal size analysis in male and female mice switched from chow to 60% HFD (A-D, J-K, N-O) for 4 weeks or 45% HFD (E-H, L-M, P-Q) for 3 weeks. Histograms of meal size by (A) grams and (B) calories in male and (C) grams and (D) calories in female mice switched to a 60% HFD. Histograms of meal size by (E) grams and (F) calories in male and (G) grams and (H) calories in female mice switched to a 45% HFD. Histogram bin sizes of 0.02 (range: 0-0.5) for meals by grams and bin size of 0.2 (range: 0-3) for meals by calories. Non-linear regression was used to fit curves on each graph. (I) Range of meal sizes in grams consumed by mice (n=16) during the chow feeding period. Quartiles were used to determine cut-offs for meal types. Type 1: <60 mg, Type2: <160 mg, Type 3: <280 mg, and Type 4: >280 mg. Pie-charts depicting meal types across diet switch week in (J) males and (K) females switched from chow to a 60% HFD and (L) males and (M) females switched from chow to a 45% HFD. Percent frequency of meals by type across each week of feeding in (N) males and (O) females switched from chow to a 60% HFD and (P) males and (Q) females switched from chow to a 45% HFD. Data are shown as mean ± SEM for N=8 per sex and were statistically analyzed using a repeated measures one-way ANOVA with Tukey multiple comparison tests. P-value definitions: not-significant (ns) >0.05, * <0.05, **<0.01, ***<0.001, **** <0.00001.

Using quartile distribution of meal sizes by mass in chow-fed male mice to yield four meal types (**Figure 5I** and **Methods**), male mice switched to 60% HFD increase the proportion of type II meals consumed while decreasing the proportion of very small type I meals and displaying a trend towards decreasing the proportion of large and very large type III and type IV meals (**Figure 5J, 5N**). This is observed across light and dark cycles (**Supplemental Figure 10A-D**). Female mice switched to 60% HFD increase the proportion of type II meals and decrease the proportion of very large type IV meals consumed, but they also increase the proportion of very small type I meals consumed (**Figure 5K, 5O**). These shifts are primarily observed during the dark cycle (**Supplemental Figure 10A-D**). Similar shifts are observed in mice switched to 45% HFD (**Figure 5L, 5M, 5P, 5Q** and **Supplemental Figure 10E-H**).

### Metabolic phenotypes and meal patterns in mice switched to 60% HFD with running wheel access

We investigated whether running wheel activity increases EE sufficiently to offset the caloric content of consuming 60% HFD. Food intake is similar in mice with active running wheels (Wheels Unlocked) and mice with inactive running wheels (Wheels Locked), with similar patterns of transient changes during the first ∼2 weeks culminating in a sustained increase in daily caloric intake (**Figure 6A-B**). EE is higher in the Wheels Unlocked group (**Figure 6C**, **Supplemental Figure 11A** for ANCOVA) due to effects during the dark cycle (**Supplemental Figure 11B**). This reduces the deviation towards positive energy balance in the Wheels Unlocked group, although net energy balance is not significantly different between groups (**Figure 6D-E, Supplemental Figure 11C**). The Wheels Unlocked group is protected against significant body weight gain during the first two weeks of 60% HFD (**Figure 6F, Supplemental Figure 11D**). However, mice with access to active running wheels begin to gain weight after the second week, and this is associated with increased food and energy intake during the dark cycle (**Supplemental Figure 12A-B**).

**Figure 6.**
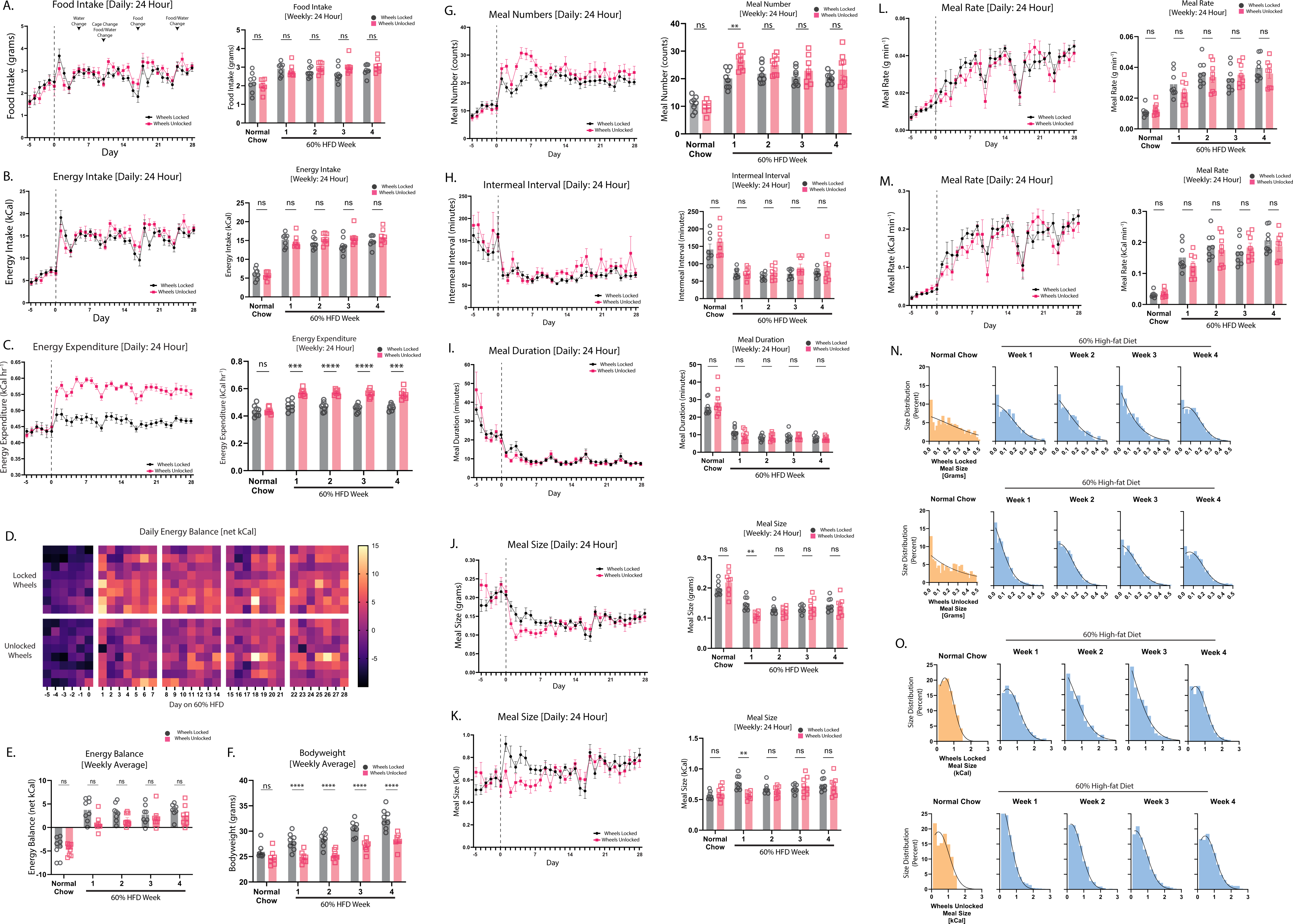
Effects of running wheel access on feeding behavior and energy expenditure in male mice switched to 60% HFD. Male mice were placed in metabolic chambers with chow and locked running wheels for 5 days. Mice were then switched to 60% HFD on day 0 (dashed line) and concomitantly remained with locked wheels (wheels locked, black) or the running wheels were unlocked (wheels unlocked, red) for 28 days. (A) Daily and average weekly food intake in grams. (B) Daily and average weekly caloric intake. (C) Daily and average weekly energy expenditure. (D) Heat map depicting net energy balance for each mouse across time after being switched from normal chow to 60% HFD with locked or unlocked running wheels. (E) Average net daily energy balance across each week of diet. (F) Average body weight across each week of diet. Feeding behavior analysis depicting (G) daily and average weekly average meal number, (H) daily and average weekly intermeal interval, (I) daily and average weekly meal duration, (J) daily and average weekly meal size in grams, (K) daily and average weekly meal size in calories, (L) daily and average weekly meal rates in grams per minute, and (M) daily and average weekly meal rate in calories per minute. (N) Histograms of meal size by gram in bin sizes of 0.02 (Range 0-0.5). (O) Histograms of meal size by calories in bin sizes of 0.2 (Range 0-3). Non-linear regression was used to fit curves on each graph. Arrows indicate external interventions such as adding food/water or cage changes. Data are shown as mean ± SEM for N=8 per group and were statistically analyzed using a two-way ANOVA with Sidaks multiple comparison tests. P-value definitions: not-significant (ns) >0.05, * <0.05, **<0.01, ***<0.001, **** <0.00001.

The Wheels Unlocked group displays a greater increase in meal number compared to the Wheels Locked group during the first week (**Figure 6G**), mainly during the dark cycle (**Supplemental Figure 12C**). Inter-meal interval and meal duration decrease equally in both groups (**Figure 6H-I, Supplemental Figure 12D-E**). Access to active running wheels lowers meal size by mass and calories on the first week of 60% HFD during both the light and dark cycles (**Figure 6J-K; Supplemental Figure 12F-G**). This is no longer evident after week 2, coincident with greater deviations towards positive energy balance and weight gain in mice with access to active running wheels. Meal rates are not significantly different between groups (**Figure 6L-M, Supplemental Figure 12H-I**). The proportion of meals consumed with a lower caloric content is higher in the Wheels Locked group (**Figure 6N-O**). Furthermore, both groups increase their consumption of type II meals and decrease their consumption of very large type IV meals, but mice with access to active running wheels also increase the consumption of very small type I meals and have a larger decrease in the proportion of very large type IV meals consumed during the first week of 60% HFD exposure (**Supplemental Figure 12J-L**).

RER is significantly lower in mice with running wheel access during the first two weeks following diet switch (**Supplemental Figure 11E-F**). Water intake decreases equally in both groups following 60% HFD exposure, although dark cycle water intake tends to be higher in the Wheels Unlocked group (**Supplemental Figure 11G-H**). Off-wheel activity decreases in both groups when switched to 60% HFD, although mice with active running wheels have higher off-wheel activity during the last two weeks of diet exposure, particularly during the dark cycle (**Supplemental Figure 11I-J**). Wheel running activity progressively increases and levels out by week four and is highest during the dark cycle (**Supplemental Figure 11K-L**).

## Discussion

Our results reveal the time course over which mice engage processes to maintain neutral energy balance and when these processes ultimately fail in the presence of HFD. Obtaining these results in the genetically tractable mouse can be leveraged towards investigating mechanisms characterizing the transition from maintaining weight homeostasis to the failure of these attempts. Furthermore, we provide analyses of other parameters, such as meal size, that we hypothesize are significant contributors to the propensity to gain weight.

Short-term (3-7 day) exposure to HFD promotes transient hyperphagia in mice (8–11). We show that transient hyperphagia followed by attempts to restore normal energy intake occurs during the first ∼14 days of HFD exposure, and mice then display sustained hyperphagia. This suggests that homeostatic mechanisms are engaged during the first ∼2 weeks of HFD exposure, but they ultimately fail or are overtaken by other factors such as hedonic mechanisms. Identifying mechanisms engaged during the first 2 weeks of HFD exposure that then fail or are overcome will provide important information about molecular targets essential for weight control. Feeding during early stages of HFD exposure is regulated by astrocyte-mediated control of brainstem glutamatergic neurotransmission from vagal inputs (11). Furthermore, the anorectic effect of cholecystokinin (CCK), a hormone regulating meal size via vagal inputs from the gut (24), is impaired in male mice after 5 weeks on HFD (17). Collectively, these findings suggest that vagal mechanisms linking the gut and brainstem are targets for future studies investigating processes contributing to weight gain in response to HFD.

Our findings agree with studies in rats showing increased meal size by caloric content upon HFD exposure (4–7). Meal size by caloric content is a strong predictor of weight gain in rats (4, 7). An increase in meal size is indicative of impaired satiation – fullness during meal ingestion. CCK-induced satiation is impaired in male mice after 5 weeks on HFD (17). Obesogenic diets also impact satiation by impairing vagal afferent responsiveness to distension and changes in gastric emptying (25, 26). Since these studies on CCK and vagal responsiveness were conducted in response to long-term HFD exposure, it is unclear whether these factors are a cause or consequence of weight gain. Our findings that meal size increases prior to weight gain strongly suggest that impaired satiation contributes to obesity. Contrasting results in rats (4, 5), we show meal number increases in mice exposed to HFD, suggesting that satiety – fullness between meals – is also impaired and highlights the need to study mechanisms that integrate satiation and satiety as contributors to weight gain.

There are interesting sex differences in the response to HFD. Male mice increase the proportion of slightly larger (type II) meals at the expense of smaller (type I) meals typically consumed during the chow period. Females also increase the proportion of type II meals consumed but partially offset this by increasing the proportion of type I meals. Male mice also display greater increases in meal rates relative to female mice. This likely contributes to the more pronounced weight gain that is typically observed in male vs. female mice. In support of this, increased energy intake and meal sizes are observed in male relative to female mice following 5 weeks on HFD (17). These sex differences can be leveraged to investigate potential mechanisms regulating meal patterns such as steroid hormone signaling (27–29).

Our observation that exposure to HFD increases energy intake during the light cycle highlights the relative importance of meal timing noted in humans and rodents. Increased light cycle energy intake is associated with greater weight gain in rodents (18, 30), and caloric consumption later in the day is associated with greater weight gain in humans (31–34). The present studies demonstrate increased caloric intake during the light cycle associated with more frequent and rapid intake of larger meals in HFD-fed mice, particularly male mice. HFD impairs vagal afferent circadian rhythms (35, 36), which may contribute to increased daytime meal size. Increasing intestinal glucagon-like peptide-1 (GLP-1) decreases daytime food intake in mice in a vagal GLP-1 receptor-mediated manner, presumably through modulating meal termination (37, 38). Thus, future studies should address whether defects in these mechanisms occur in a time scale corresponding to the changes in feeding behavior we demonstrate in the present studies.

Male and female mice experience quantitatively similar increases in absolute EE when switched to HFD, but this is not sufficient to compensate for the increased caloric intake. Protection from weight gain in female relative to male mice may be driven by the ability of female mice to better utilize fat (9, 17), supported by our data showing lower relative RER in female compared to male mice switched to a 60% HFD. This also agrees with recent studies showing that female mice balance energy expenditure and intake more efficiently than do male mice (39) EE increases in response to HFD even as locomotor activity decreases, as previously observed (10, 40). Decreased locomotor activity in HFD-fed mice accounts for >60% of weight gain (40). Providing male mice with access to running wheels when switched to 60% HFD significantly increases EE and partially mitigates the shortfall between EE and caloric intake, thus reducing weight gain, as previously shown (41). Satiation is also enhanced in mice with wheel running access, and we speculate this further contributes to reduced weight gain. However, after ∼2 weeks of HFD/running wheel exposure, enhanced satiation is no longer apparent and mice with wheel running access begin to gain weight. These findings further emphasize the potential role of meal size as a contributor to weight gain in the presence of obesogenic diets.

In sum, our study provides insights into how time-dependent changes in feeding behaviors following obesogenic diet exposure can modify weight gain. Although previous studies in rats show similar findings, the present studies conducted in the more genetically tractable mouse provide the foundation for future studies to investigate molecular mechanisms relevant to the regulation of meal patterns and weight gain. Furthermore, our analyses using different HFD and comparing sexes not only further emphasize the importance of the association between meal patterns and weight gain but also highlight phenotypes that can be used in future studies on the contribution of diet composition and sex on weight gain.

### Considerations

Our studies were conducted at a standard housing temperature for most animal facilities (23°C), which is below the thermoneutral temperature for mice (∼28-30°C). Temperature impacts EE and weight gain in a sex-dependent manner in mice (39), so repeating these experiments at a higher temperature will likely yield different outcomes regarding EE and meal patterns. The present studies did not use matched low-fat diets for each HFD. The magnitude of metabolic changes may be different when using matched low-fat diets. Although we established visual and threshold parameters for food spillage (see **Methods**), there is inherent inaccuracy in food intake measurements resulting from food spillage and caching occurring with any metabolic chamber. We did not measure fecal energy content, so the energy balance calculation assumes complete energy absorption. Finally, interpreting running wheel results must consider the fact that mice were provided with two rewarding stimuli simultaneously (HFD and running). Our colleague Dr. Nathan Winn shows that switching mice to a 60% HFD after 5 weeks of running wheel access further increases running wheel activity, but increased caloric intake is equivalent to mice switched to 60% HFD without prior access to running wheels (personal communication).

## Author Contributions

Conceptualization: JEA, MBB, PF; Methodology: JEA, MBB, PF; Analysis: JEA, PF; Writing: JEA, PF; Funding: JEA. All authors reviewed and have approved the final version of this manuscript.

## Acknowledgements and Funding

We would like to thank Dr. Louise Lantier and Merrygay James for their assistance. Promethion metabolic chamber studies were conducted within the Vanderbilt Mouse Metabolic Phenotyping Center (VMMPC) which is supported by NIH grant U2C0595637, shared instrument grant S10OD028455 (JEA). This work was funded by NIH grants F31-DK127728 (PF), T32-DK07563 (PF, MBB), and R01-DK132852 (JEA).

---

**Supplemental Figure 1.**
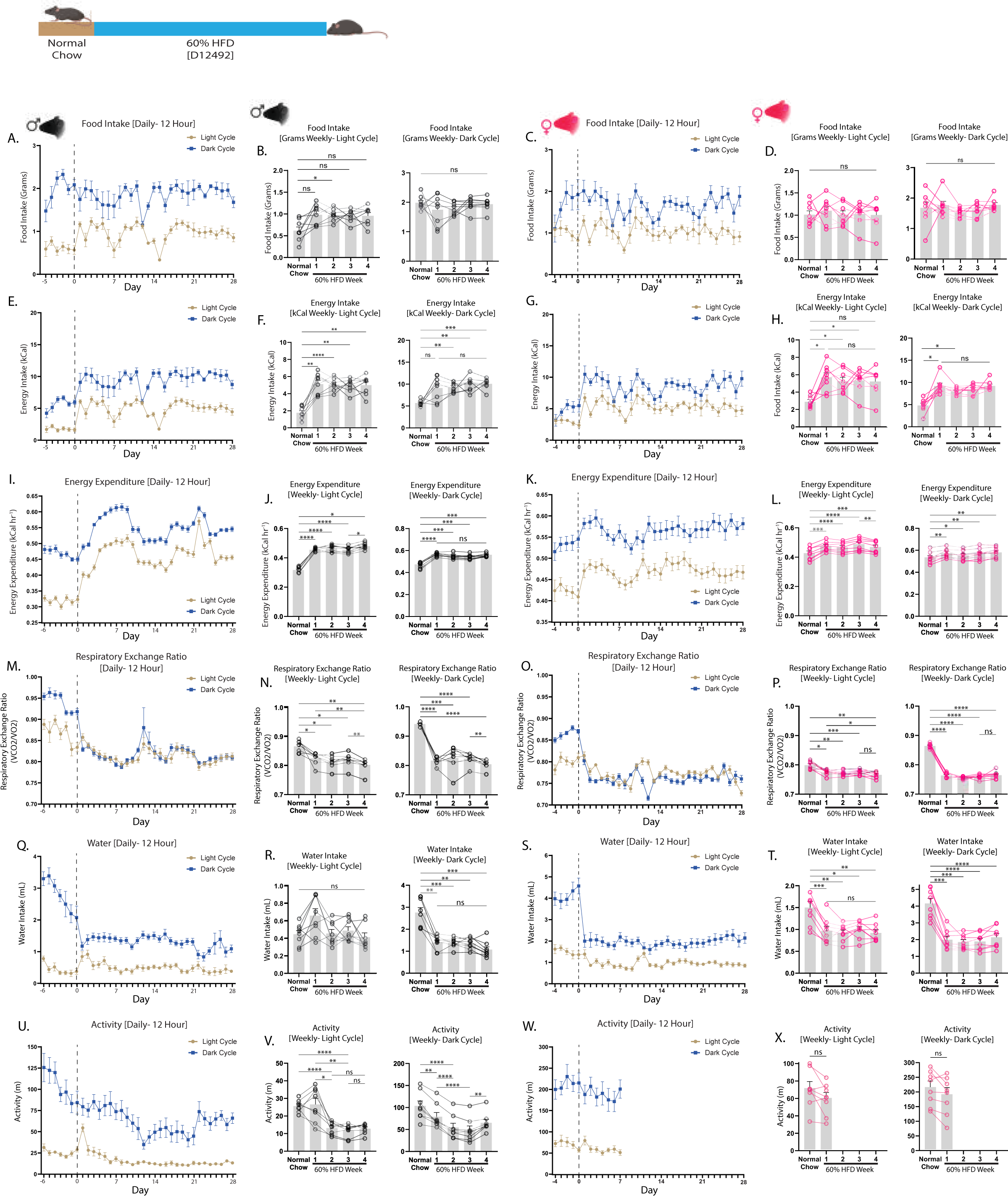
Metabolic phenotyping of mice switched to 60% HFD separated by light-dark cycle. Metabolic phenotyping parameters in male and female mice switched from chow to 60% HFD on day 0 (dashed lines) in metabolic chambers and divided by light (blue) and dark (beige) cycles. (A) Daily food intake in grams in male and (C) female mice. Average weekly food intake by mass in the light and dark cycle in (B) male mice and in (D) female mice. (E) Daily caloric intake in male and (G) female mice. Average weekly caloric intake in the light and dark cycle in (F) male mice and in (H) female mice. (I) Daily energy expenditure in male and (K) female mice. Average weekly energy expenditure in the light and dark cycle in (J) male mice and in (L) female mice. (M) Daily respiratory exchange ratio in male and (O) female mice. Average weekly respiratory exchange ratio in the light and dark cycle in (N) male mice and in (P) female mice. (Q) Daily water intake in male and (S) female mice. Average weekly water intake in the light and dark cycle in (R) male mice and in (T) female mice. (U) Daily locomotor activity in male and (W) female mice. Average weekly locomotor activity in the light and dark cycle in (V) male mice and in (X) female mice. Locomotor activity data are only available for first seven days of diet switch in female mice due to instrument malfunction for this feature. Data are shown as mean ± SEM for N=8 per sex and were statistically analyzed using a repeated measures one-way ANOVA with Tukey multiple comparison tests. P-value definitions: not-significant (ns) >0.05, * <0.05, **<0.01, ***<0.001, **** <0.00001.

**Supplemental Figure 2.**
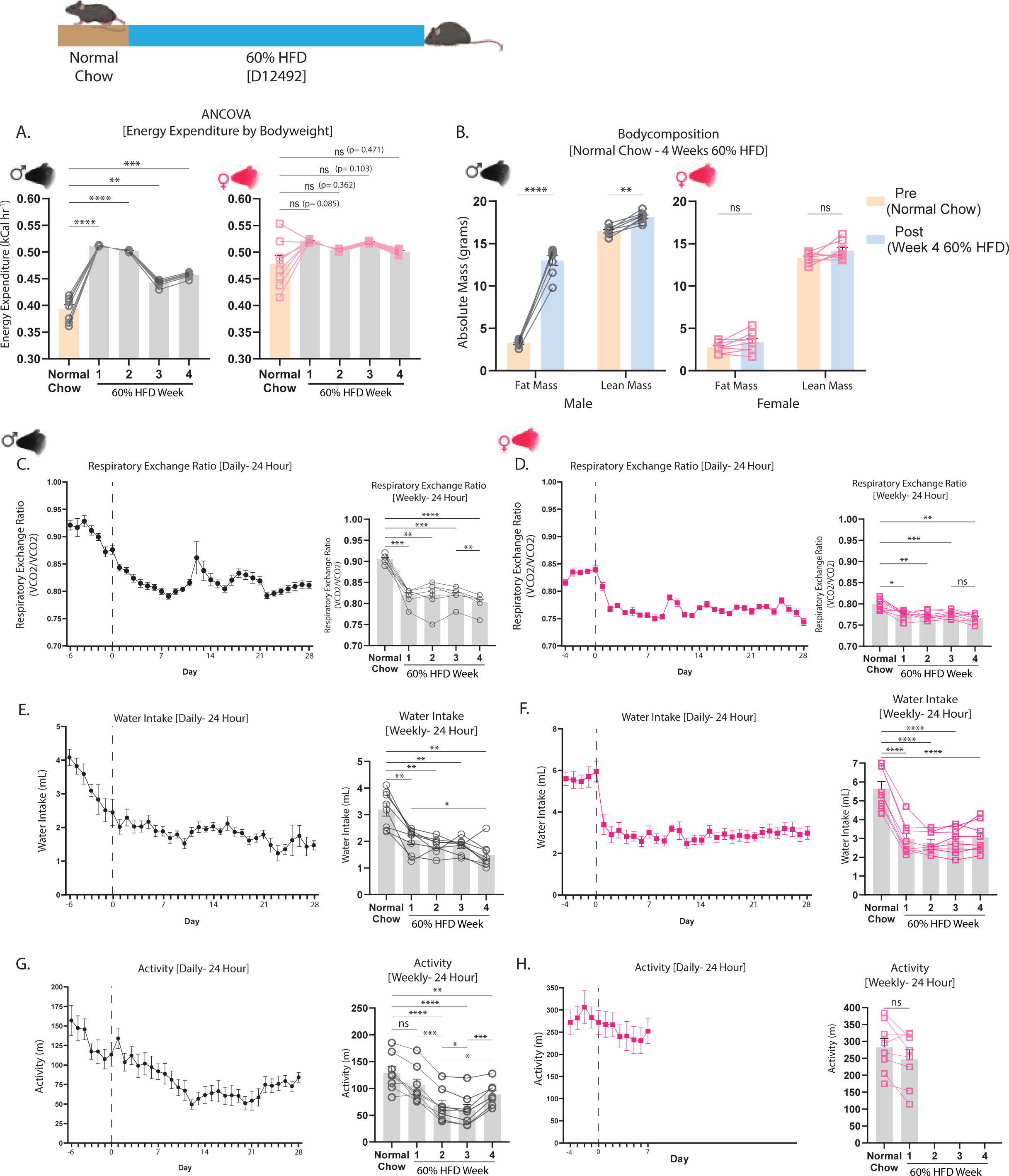
Metabolic phenotyping of mice switched to 60% HFD. Metabolic parameters in male (black) and female (pink) mice fed a chow diet and switched to 60% HFD for 28 days. (A) ANCOVA of average weekly energy expenditure with body weight as a covariate. (B) Body composition in chow-fed (Pre) and at the end of 28 days of HFD feeding (Post). (C) Daily and average weekly respiratory exchange ratio in male and (D) female mice. (E) Daily and average weekly water intake in male and (F) female mice. (G) Daily and average weekly locomotor activity in male and (H) female mice. Locomotor activity data are only available for first seven days of diet switch in female mice due to instrument malfunction for this feature. Data are shown as mean ± SEM for N=8 per sex and were statistically analyzed using a repeated measures one-way ANOVA with Tukey multiple comparison tests. P-value definitions: not-significant (ns) >0.05, * <0.05, **<0.01, ***<0.001, **** <0.00001.

**Supplemental Figure 3.**
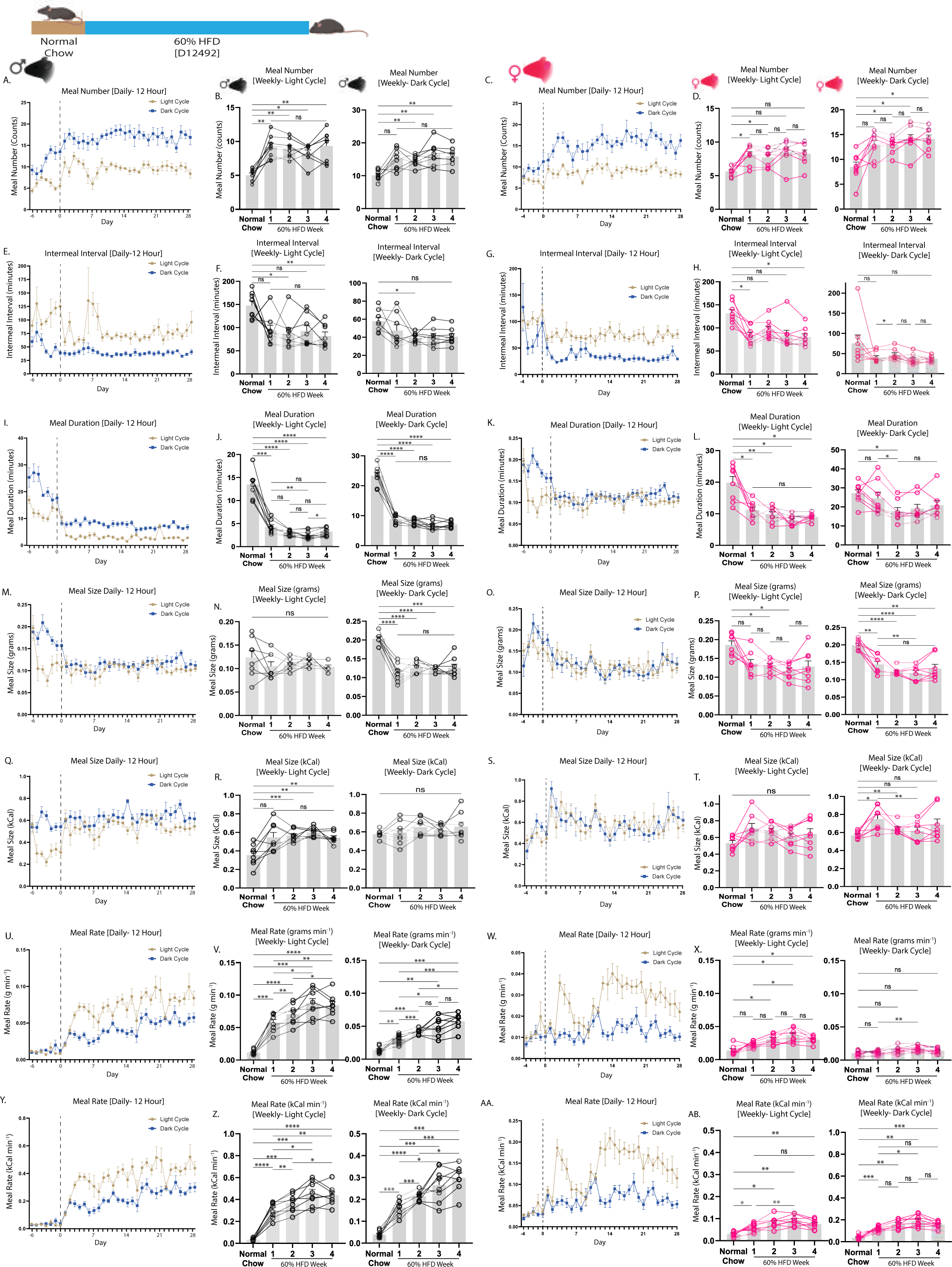
Feeding behaviors in mice switched to 60% HFD separated by light-dark cycle. Feeding behavior analysis in male and female mice switched from chow to 60% HFD on day 0 (dashed lines) in metabolic chambers and divided by light (blue) and dark (beige) cycles. (A) Daily and (B) average weekly meal number in male and (C) daily and (D) average weekly meal number in female mice. (E) Daily and (F) average weekly inter-meal interval in male mice and (G) daily and (H) average weekly inter-meal interval in female mice. (I) Daily and (J) average weekly meal duration in male mice and (K) daily and (L) average weekly meal duration in female mice. (M) Daily and (N) average weekly meal size by mass in male mice and (O) daily and (P) average weekly meal size by mass in female mice. (Q) Daily and (R) average weekly meal size by calorie in male mice and (S) daily and (T) average weekly meal size by calorie in female mice. (U) Daily and (V) average weekly meal rate in grams per minute in male mice and (W) daily and (X) average weekly meal rate in grams per minute in female mice. (Y) Daily and (Z) average weekly meal rate in calories per minute in male mice and (AA) Daily and (AB) average weekly meal rate in calories per minute in female mice. Data are shown as mean ± SEM for N=8 per sex and were statistically analyzed using repeated measures one-way ANOVA with Tukey multiple comparison tests. P-value definitions: not-significant (ns) >0.05, * <0.05, **<0.01, ***<0.001, **** <0.00001.

**Supplemental Figure 4.**
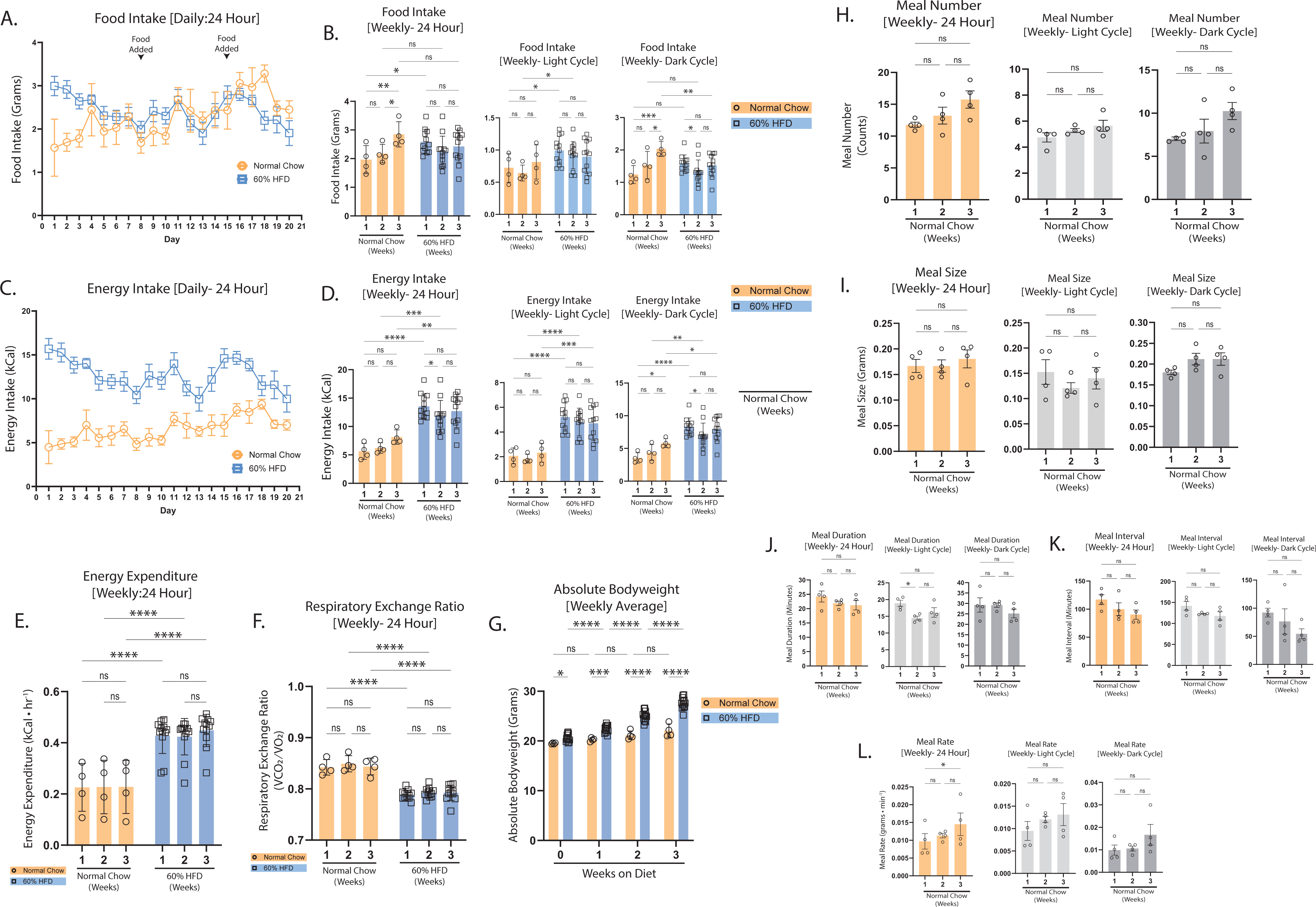
Food intake, energy expenditure, and feeding behaviors in male mice retained on chow or switched to 60% HFD. Male mice were switched to 60% HFD on day 0 or remained on chow for 21 days. (A) Daily food intake by mass and (B) average weekly food intake by mass over 24h and during the light and dark cycle. (C) Daily energy intake by calorie and (D) average weekly energy intake by calorie over 24h and during the light and dark cycle. (E) Daily energy expenditure and (F) average weekly energy expenditure over 24h and during the light and dark cycle. (G) Average weekly body weight. (H) Average weekly meal number by 24h, light cycle, and dark cycle in chow-fed mice. (I) Average meal size by mass by 24h, light cycle, and dark cycle in chow-fed mice. (J) Average meal duration by 24h, light cycle, and dark cycle in chow-fed mice. (K) Average inter-meal interval by 24h, light cycle, and dark cycle in chow-fed mice. (L) Average weekly meal rate by mass by 24h, light cycle, and dark cycle in chow-fed mice. Data are shown as mean ± SEM for N=4-12 per group and were statistically analyzed using repeated measures one-way ANOVA with Tukey multiple comparison tests or two-way ANOVA with Sidak multiple comparison test. P-value definitions: not-significant (ns) >0.05, * <0.05, **<0.01, ***<0.001, **** <0.00001.

**Supplemental Figure 5.**
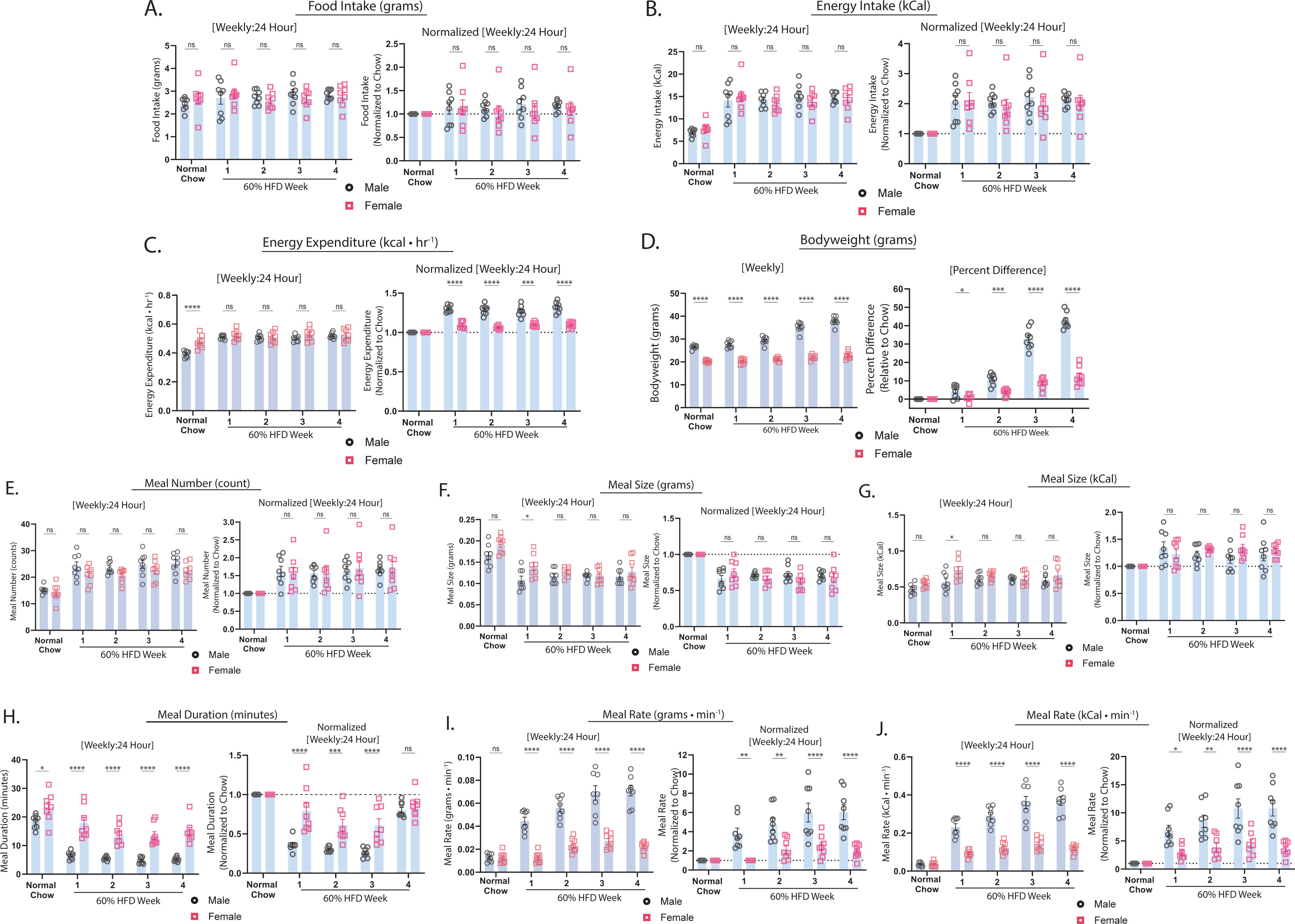
Effects of sex on 60% HFD-induced changes in metabolic parameters and meal patterns. Average weekly data are shown for male (black symbols) and female (pink symbols) mice as both absolute values and values normalized to chow for each individual mouse across 24-hour cycles. (A) Food intake by mass. (B) Energy intake by calorie. (C) Energy expenditure. (D) Body weight. (E) Meal number. (F) meal size by mass. (G) Meal size by calorie. (H) Meal duration. (I) Meal rate by mass. (J) Meal rate by calorie. Data are shown as mean ± SEM for N=8 per sex and were statistically analyzed using two-way ANOVA with Sidak multiple comparison test. P-value definitions: not-significant (ns) >0.05, * <0.05, **<0.01, ***<0.001, **** <0.00001.

**Supplemental Figure 6.**
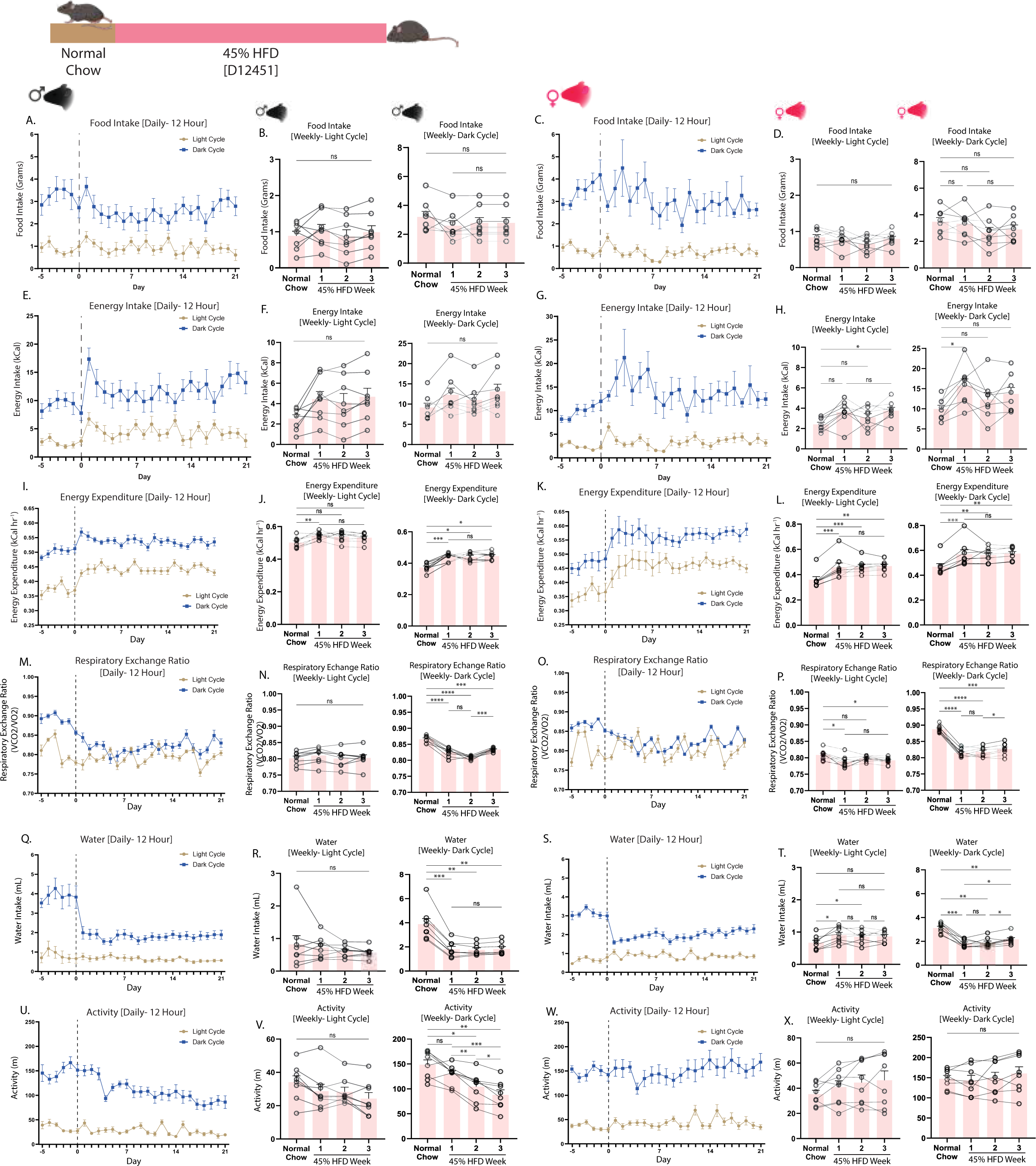
Metabolic phenotyping of mice switched to 45% HFD separated by light-dark cycle. Metabolic phenotyping parameters in male and female mice switched from chow to 45% HFD on day 0 (dashed lines) in metabolic chambers and divided by light (blue) and dark (beige) cycles. (A) Daily food intake in grams in male and (C) female mice. Average weekly food intake by mass in the light and dark cycle in (B) male mice and in (D) female mice. (E) Daily caloric intake in male and (G) female mice. Average weekly caloric intake in the light and dark cycle in (F) male mice and in (H) female mice. (I) Daily energy expenditure in male and (K) female mice. Average weekly energy expenditure in the light and dark cycle in (J) male mice and in (L) female mice. (M) Daily respiratory exchange ratio in male and (O) female mice. Average weekly respiratory exchange ratio in the light and dark cycle in (N) male mice and in (P) female mice. (Q) Daily water intake in male and (S) female mice. Average weekly water intake in the light and dark cycle in (R) male mice and in (T) female mice. (U) Daily locomotor activity in male and (W) female mice. Average weekly locomotor activity in the light and dark cycle in (V) male mice and in (X) female mice. Data are shown as mean ± SEM for N=8 per sex and were statistically analyzed using a repeated measures one-way ANOVA with Tukey multiple comparison tests. P-value definitions: not-significant (ns) >0.05, * <0.05, **<0.01, ***<0.001, **** <0.00001.

**Supplemental Figure 7.**
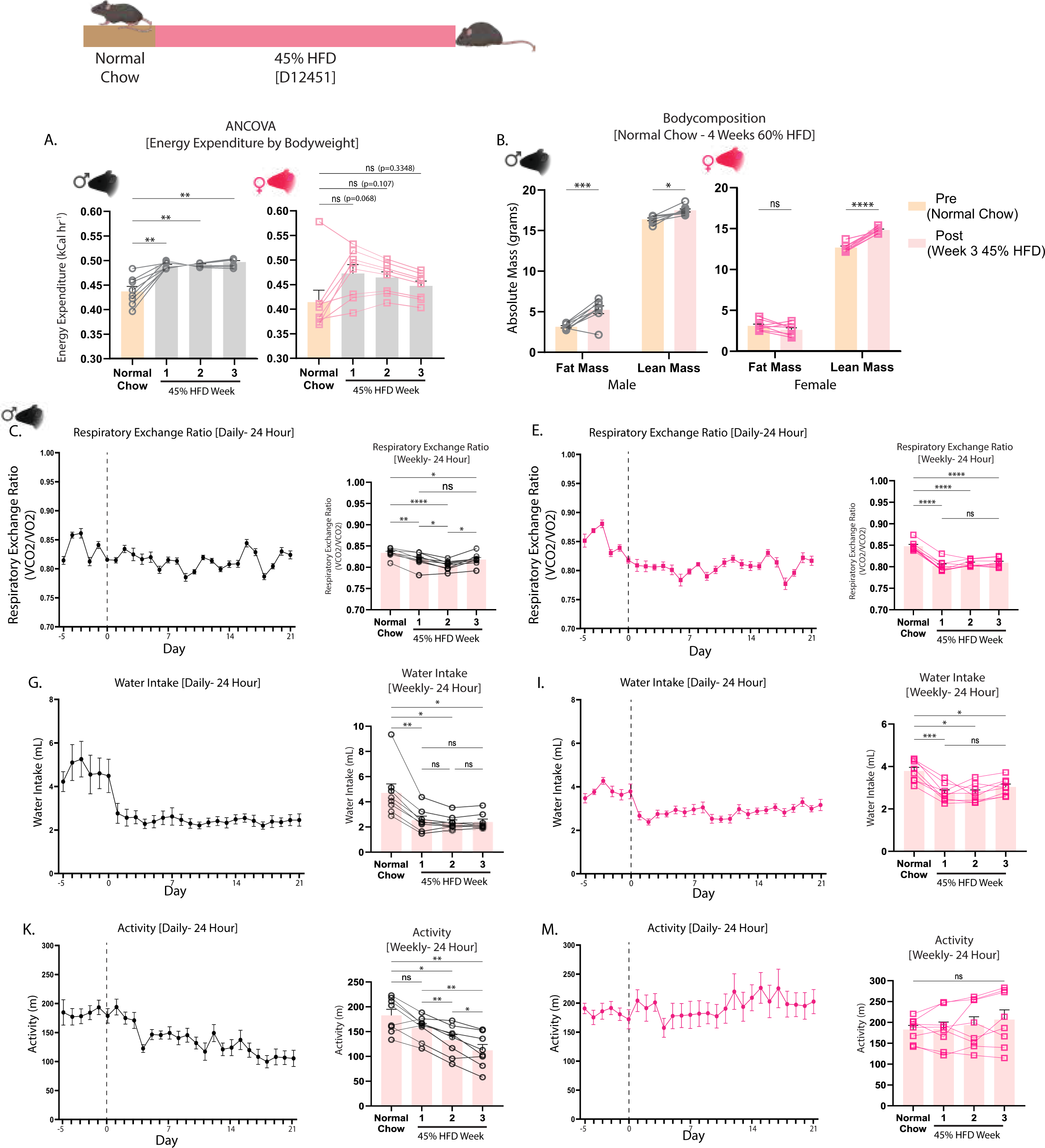
Metabolic phenotyping of mice switched to 45% HFD. Metabolic parameters in male (black) and female (pink) mice fed a chow diet and switched to 45% HFD for 21 days. (A) ANCOVA of average weekly energy expenditure with body weight as a covariate. (B) Body composition in chow-fed (Pre) and at the end of 21 days of HFD feeding (Post). (C) Daily and average weekly respiratory exchange ratio in male and (D) female mice. (E) Daily and average weekly water intake in male and (F) female mice. (G) Daily and average weekly locomotor activity in male and (H) female mice. Data are shown as mean ± SEM for N=8 per sex and were statistically analyzed using a repeated measures one-way ANOVA with Tukey multiple comparison tests. P-value definitions: not-significant (ns) >0.05, * <0.05, **<0.01, ***<0.001, **** <0.00001.

**Supplemental Figure 8.**
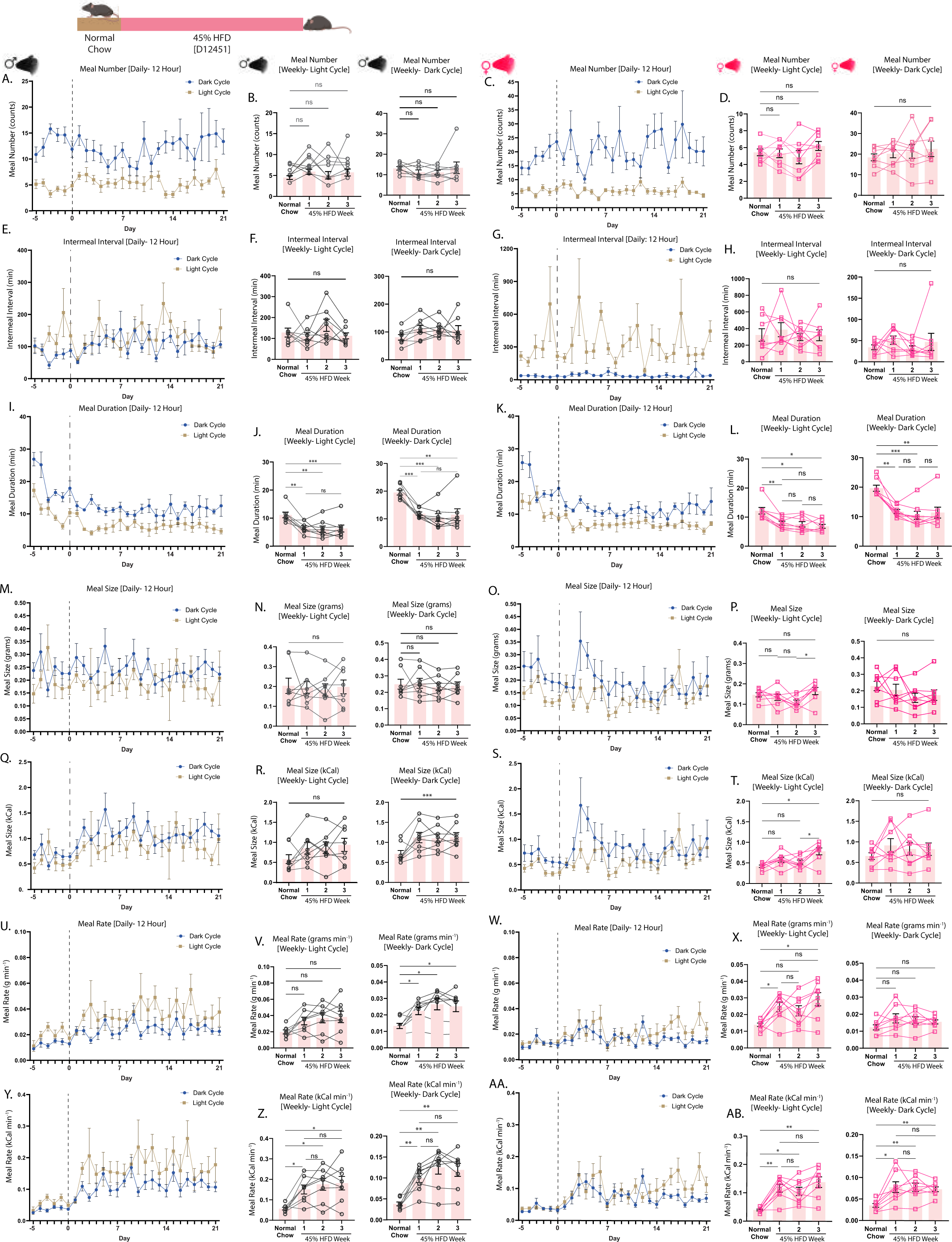
Feeding behaviors in mice switched to 45% HFD separated by light-dark cycle. Feeding behavior analysis in male and female mice switched from chow to 45% HFD on day 0 (dashed lines) in metabolic chambers and divided by light (blue) and dark (beige) cycles. (A) Daily and (B) average weekly meal number in male and (C) daily and (D) average weekly meal number in female mice. (E) Daily and (F) average weekly inter-meal interval in male mice and (G) daily and (H) average weekly inter-meal interval in female mice. (I) Daily and (J) average weekly meal duration in male mice and (K) daily and (L) average weekly meal duration in female mice. (M) Daily and (N) average weekly meal size by mass in male mice and (O) daily and (P) average weekly meal size by mass in female mice. (Q) Daily and (R) average weekly meal size by calorie in male mice and (S) daily and (T) average weekly meal size by calorie in female mice. (U) Daily and (V) average weekly meal rate in grams per minute in male mice and (W) daily and (X) average weekly meal rate in grams per minute in female mice. (Y) Daily and (Z) average weekly meal rate in calories per minute in male mice and (AA) Daily and (AB) average weekly meal rate in calories per minute in female mice. Data are shown as mean ± SEM for N=8 per sex and were statistically analyzed using repeated measures one-way ANOVA with Tukey multiple comparison tests. P-value definitions: not-significant (ns) >0.05, * <0.05, **<0.01, ***<0.001, **** <0.00001.

**Supplemental Figure 9.**
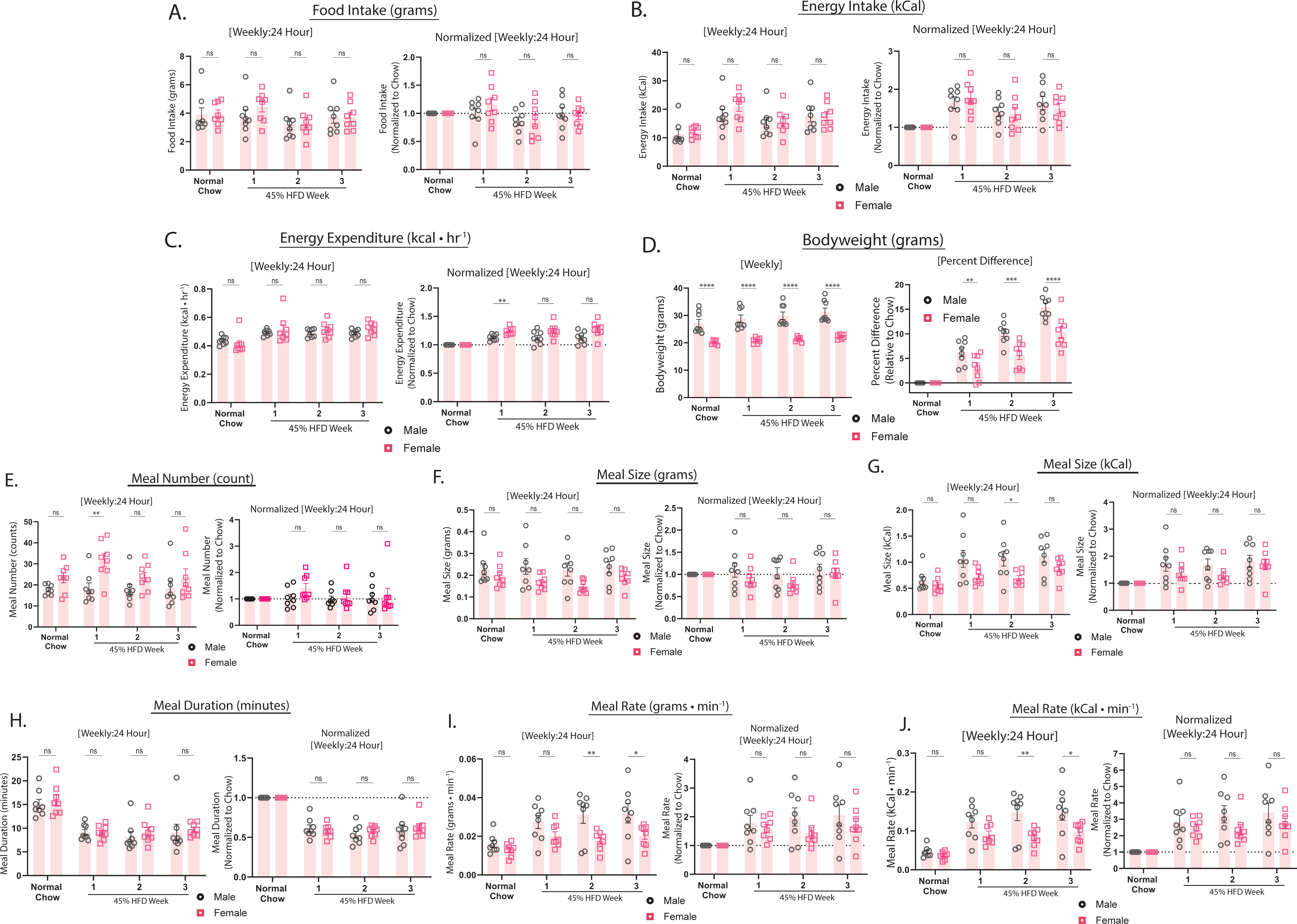
Effects of sex on 45% HFD-induced changes in metabolic parameters and meal patterns. Average weekly data are shown for male (black symbols) and female (pink symbols) mice as both absolute values and values normalized to chow for each individual mouse across 24-hour cycles. (A) Food intake by mass. (B) Energy intake by calorie. (C) Energy expenditure. (D) Body weight. (E) Meal number. (F) Meal size by mass. (G) Meal size by calorie. (H) Meal duration. (I) Meal rate by mass. (J) Meal rate by calorie. Data are shown as mean ± SEM for N=8 per sex and were statistically analyzed using two-way ANOVA with Sidak multiple comparison test. P-value definitions: not-significant (ns) >0.05, * <0.05, **<0.01, ***<0.001, **** <0.00001.

**Supplemental Figure 10.**
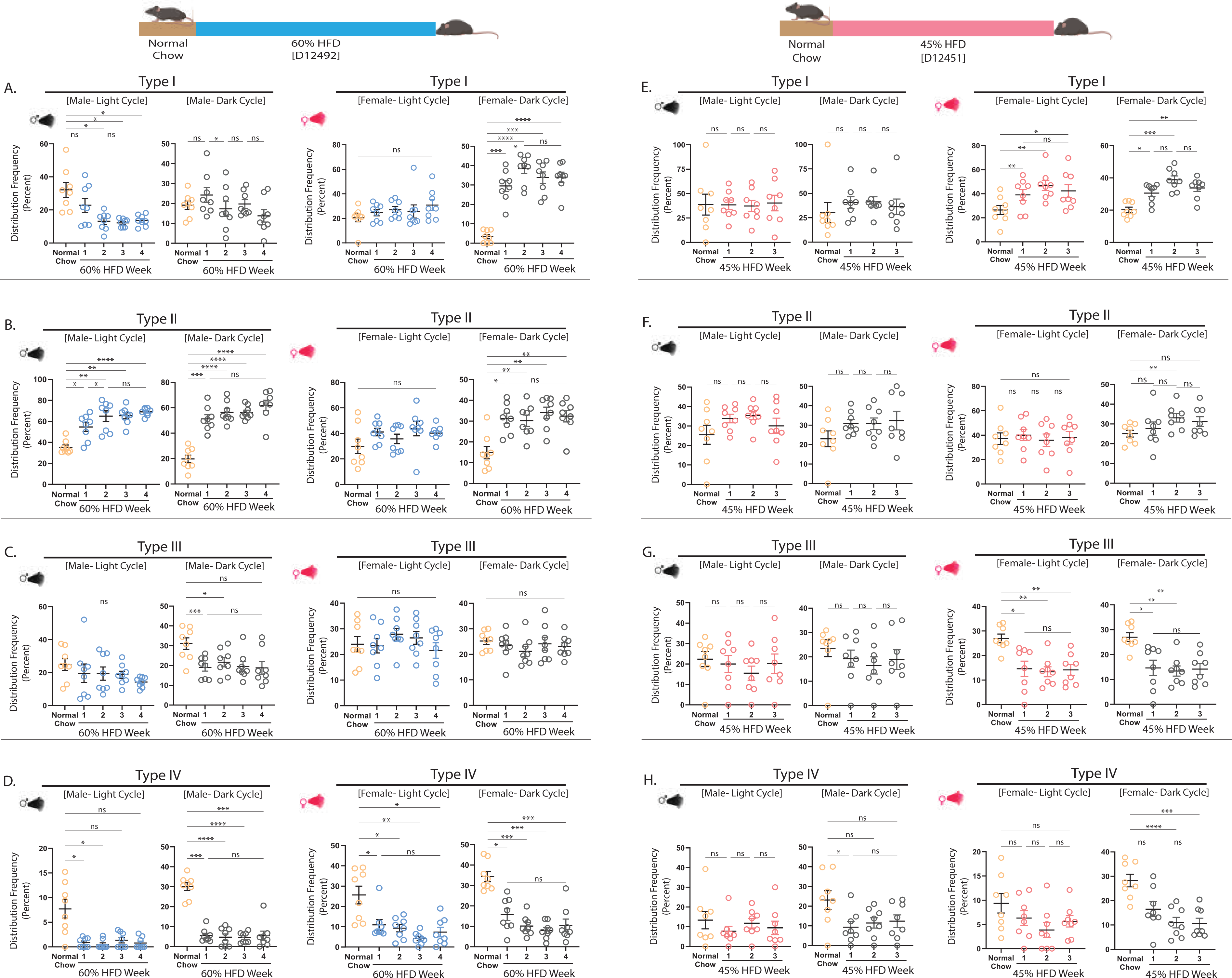
Shifts in meal patterns across time and light-cycle in mice fed obesogenic diets. Frequency distribution of Type I-IV meals in male and female mice fed chow and then switched to 60% HFD (A-D) or 45% HFD (E-H). (A, E) Type I (<60 mg), (B, F) Type II (<160 mg), (C, G) Type III (<280 mg), (D, H) Type IV (>280 mg) meals. Data are shown as mean ± SEM for N=8 per sex and were statistically analyzed using a repeated measures one-way ANOVA with Tukey multiple comparison tests. P-value definitions: not-significant (ns) >0.05, * <0.05, **<0.01, ***<0.001, **** <0.00001.

**Supplemental Figure 11.**
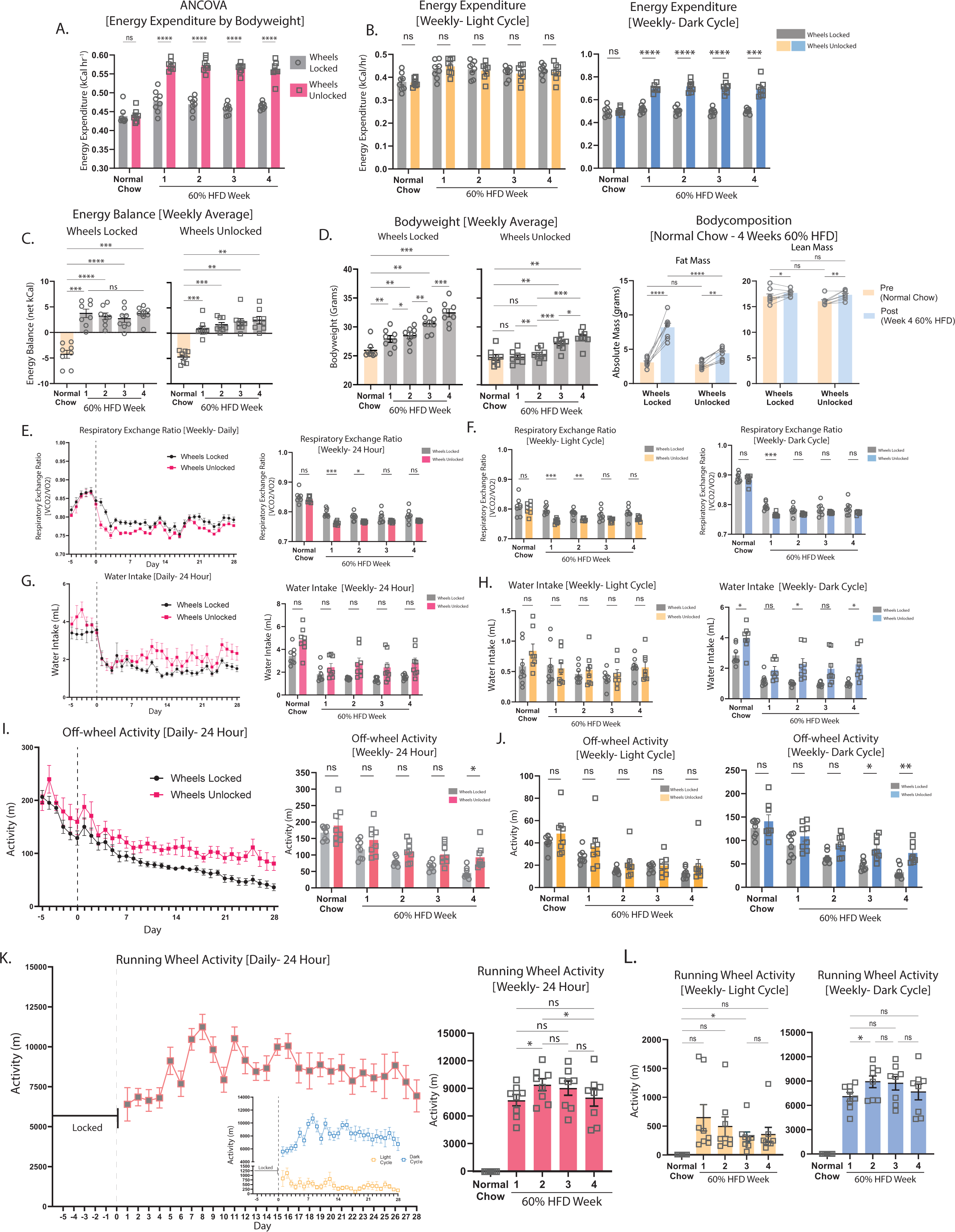
Metabolic phenotyping of mice switched to 60% HFD with running wheel access. Metabolic parameters in male mice switched from chow to a 60% HFD at day 0 (dashed line) in metabolic chambers with wheels locked or wheels unlocked. (A) ANCOVA of average weekly energy expenditure with body weight as a covariate. (B) Average weekly energy expenditure during the light and dark cycles. (C) Average weekly energy balance. (D) Average weekly body weight and body composition. (E) Daily and average weekly respiratory exchange ratio and (F) average weekly respiratory exchange ratio during the light and dark cycle. (G) Daily and average weekly water intake and (H) average weekly water intake during the light and dark cycle. (I) Daily and average weekly off-wheel activity and (J) average weekly off-wheel activity during the light and dark cycle. (K) Daily and average weekly running wheel activity and (L) average weekly running wheel activity during the light cycle and dark cycle in mice with access to unlocked running wheels. Data are shown as mean ± SEM for N=8 per group and were statistically analyzed using a two-way ANOVA with Sidaks multiple comparison tests. P-value definitions: not-significant (ns) >0.05, * <0.05, **<0.01, ***<0.001, **** <0.00001.

**Supplemental Figure 12.**
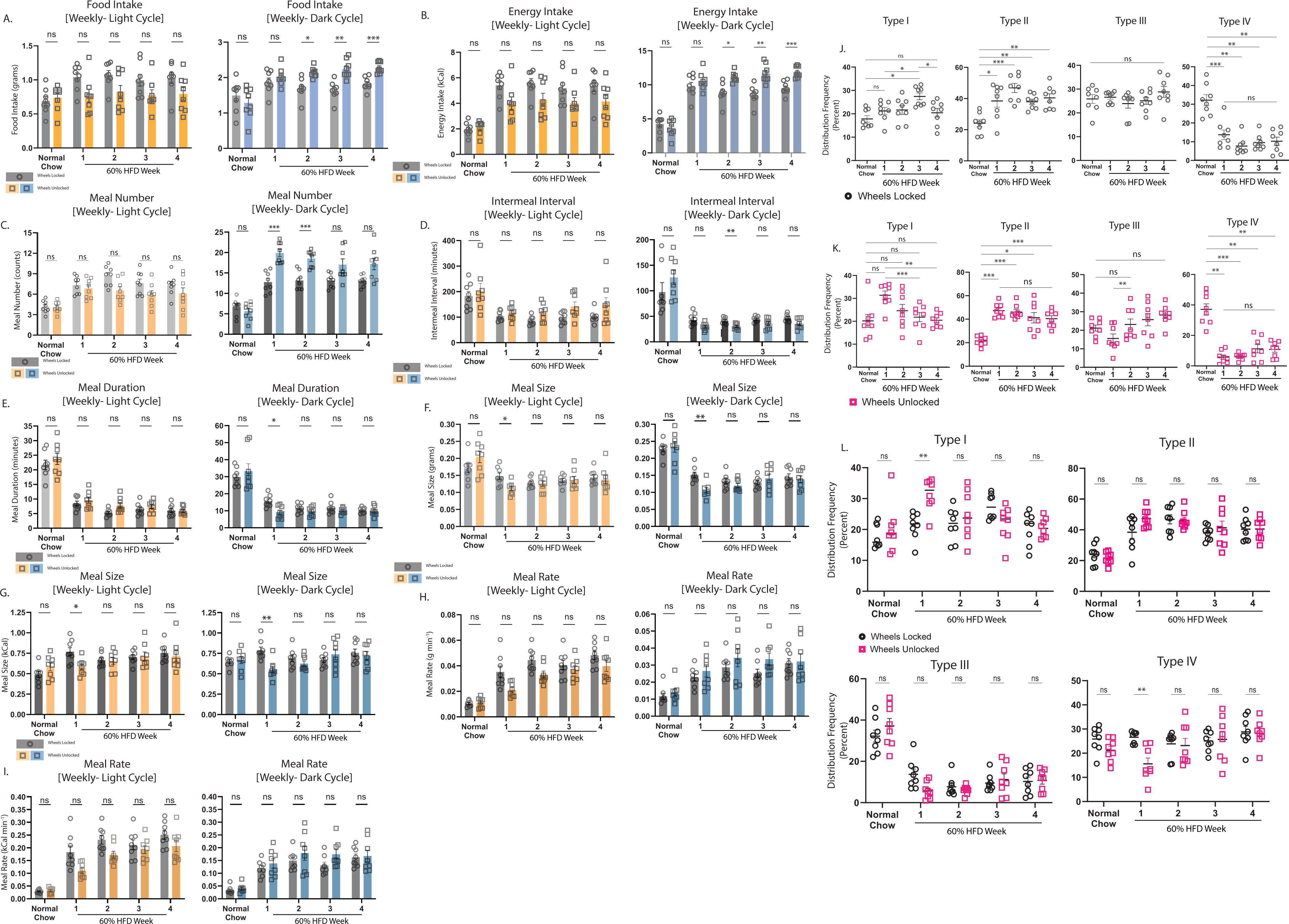
Effects of running wheel access on feeding behaviors and meal patterns. Feeding behavior and meal type analysis by light-dark cycle in male mice switched from chow to a 60% HFD with access to locked or unlocked running wheels. Average weekly (A) food intake by mass, (B) energy intake by calorie, (C) meal numbers, (D) inter-meal interval, (E) meal duration, (F) meal size by mass (G) meal size by calorie, (H) meal rates by grams per minute, and (I) meal rates by calories per minute. Mice with locked wheels are shown in light gray (light cycle) and dark gray (dark cycle) bars with data points depicted by circles. Mice with unlocked wheels are shown in orange (light cycle) and blue (dark cycle) with data points depicted by squares. Frequency of meal types [Type I (<60 mg), Type II (<160 mg), Type III (<280 mg), Type IV (>280] across time on diet in mice with (J) wheels locked and (K) wheels unlocked. (L) Frequency of meal types across time on diet in mice with wheels locked vs. wheels unlocked. Data are shown as mean ± SEM for N=8 per group and were statistically analyzed using a two-way ANOVA with Sidaks multiple comparison tests. P-value definitions: not-significant (ns) >0.05, * <0.05, **<0.01, ***<0.001, **** <0.00001.

